# SF3B1 mutant-induced missplicing of MAP3K7 causes anemia in myelodysplastic syndromes

**DOI:** 10.1101/2020.06.24.169052

**Authors:** Yen K. Lieu, Zhaoqi Liu, Abdullah M. Ali, Xin Wei, Alex Penson, Jian Zhang, Xiuli An, Raul Rabadan, Azra Raza, James L. Manley, Siddhartha Mukherjee

**Author notes:** These authors contributed equally (Zhaoqi Liu, Abdullah M. Ali). Senior author (James L. Manley, Siddhartha Mukherjee).

## Abstract

SF3B1 is the most frequently mutated RNA splicing factor in multiple neoplasms, including ~25% of myelodysplastic syndromes (MDS) patients. Mortality in MDS frequently results from severe anemia, but the underlying mechanism is largely unknown. Here we elucidate the detailed, elusive pathway by which *SF3B1* mutations cause anemia. We demonstrate, in CRISPR-edited cell models, normal human primary cells, and MDS patient cells, that mutant SF3B1 induces a splicing error in transcripts encoding the kinase MAP3K7, resulting in reduced MAP3K7 protein levels and deactivation of downstream target p38 MAPK. We show that disruption of this MAP3K7-p38 MAPK pathway leads to premature downregulation of GATA1, a master regulator of erythroid differentiation, and that this is sufficient to trigger accelerated differentiation and apoptosis. As a result, the overproduced, late staged erythroblasts undergo apoptosis and are unable to mature in the bone marrow. Our findings provide a detailed mechanism explaining the origins of anemia in MDS patients harboring *SF3B1* mutations.

## INTRODUCTION

Myelodysplastic syndromes (MDS) are a heterogeneous group of blood malignancies that originate from hematopoietic stem cells and are characterized by ineffective hematopoiesis, dysplasia, peripheral blood cytopenias, and an increased risk of transformation to acute myeloid leukemia^1,2^. Though MDS is the most common malignancy in the elderly, few effective therapies are available^3^. In the past decade, next-generation sequencing technology revealed that mutations in a number of genes that encode mRNA splicing factors constitute a common cause of MDS^4^. Among these, the most frequently mutated gene is *SF3B1*, with a frequency of 20-29% in all MDS cases and 65-83% in the subtype Refractory Anemia with Ring Sideroblasts (RARS) that is associated with red blood cell dysfunction^4-6^. Heterozygous *SF3B1* missense mutations in MDS are characterized by accumulation of erythroid precursors in the bone marrow, are significantly correlated with ringed sideroblastic anemia^5,7,8^, and by most studies, are associated with good prognosis^5,6,8,9^. *SF3B1* mutations have subsequently been found in a significant number of other cancers^6,10-12^.

SF3B1 is a subunit of the SF3B complex, which associates with the U2 snRNP during splicing and facilitates interaction of the snRNP with the pre-mRNA branch site. Several studies have reported that *SF3B1* mutations cause aberrant recognition and selection of the branch site adenosine, leading to widespread usage of cryptic 3’ splice sites (3’ss) that are approximately 10-25 nt upstream of the canonical 3′ss^13-16^. In addition, we recently provided insight into the mechanism by which *SF3B1* mutations lead to missplicing, by showing that mutant SF3B1 is unable to interact with another splicing factor, SUGP1, that itself is required for proper branchsite recognition of mutant SF3B1-sensitive introns^13^.

In an effort to elucidate the functions of mutant SF3B1, knock-in mice carrying the most common hotspot mutation in MDS, K700E, were generated^17,18^. These mice displayed erythroid dysplasia, but were not able to recapitulate other cardinal features of MDS, particularly RARS phenotypes such as accumulation of erythroid precursors in the bone marrow, increased apoptotic bone marrow cells or erythroid precursors, and appearance of ringed sideroblasts^5,7,8,19-22^. In addition, there was very little overlap of misspliced transcripts between MDS patients and the mutant mice (~ 5-10%, ^17^; ^18^), most likely due to the poor conservation of intronic sequences (~30%) between human and mouse^23^ and significant differences in alternative splicing^24^. Thus, the very little overlap of misspliced gene transcripts between MDS patients and mutant mice questions whether the observed erythroid dysplasia is a mouse-specific effect. Hence, while mutant mouse models can be valuable, they may be less useful in modeling splicing-related diseases such as MDS.

Numerous aberrantly spliced transcripts have been identified in SF3B1 mutant samples obtained from MDS patients. However, very few of these have been shown to contribute to MDS phenotypes. Recently, it was reported that missplicing of the transcript encoding the erythroid hormone erythroferrone, and the resultant decreased levels of the hormone, might be responsible for the systemic iron loading seen in MDS patients with *SF3B1* mutations who had not received blood transfusions^25^. In addition, missplicing of the transcript encoding the mitochondrial ATP-binding cassette iron transporter ABCB7 induced by mutant SF3B1 may be responsible for the accumulation of ring sideroblasts in MDS^26,27^. Despite these important insights into the identity of key mutant SF3B1 target gene transcripts, the mechanism by which *SF3B1* mutations induce anemia in MDS patients is unknown.

In this report, we provide a functional explanation for how *SF3B1* mutations cause anemia. We demonstrate, in both cell culture models and MDS patient cells, that an SF3B1 K700E-induced splicing error in transcripts encoding the MAP kinase MAP3K7, which had previously been reported to be misspliced in MDS SF3B1 mutant cells^13,26,28^, results in reduced levels of the protein and deactivation of the downstream p38 MAPK (p38). p38 is a known regulator of various biological processes such as cell differentiation and apoptosis^29^, including a pathway central to erythrocyte differentiation^30-32^. We show that disruption of this pathway leads to premature downregulation of GATA1, a master regulator of erythroid differentiation, and that this triggers accelerated erythroid differentiation and apoptosis. Thus, abnormally excessive late staged erythroblasts undergoing apoptosis are not able to mature in the bone marrow, providing a mechanistic explanation for the anemia seen in MDS patients harboring *SF3B1* mutations.

## RESULTS

### K562/SF3B1 K700E cells exhibit accelerated differentiation and increased erythroid cell death as compared to WT cells

Severe anemia is a frequent manifestation of MDS, and as such contributes to mortality. To begin to investigate how *SF3B1* mutations might lead to anemia, or other features of MDS, we used CRISPR/Cas9 technology to introduce the most common hotspot mutation found in MDS patients, K700E, into human erythroleukemia K562 cells (Fig. 1A). K562 cells were selected for their ability to differentiate along specific myeloid, megakaryocytic or erythroid lineages upon treatment with the appropriate inducer^33,34^. To avoid possible clonal or off-target effects, we isolated nine independent K700E mutant and nine independent wild-type (WT) clones, and used all of these in the experiments described below. SF3B1 mutant K700E mRNA (Fig. S1A) and protein (Fig. S1B) were stably expressed as measured by DNA sequencing of RT-PCR products and western blot (WB) analysis, respectively. Total SF3B1 protein levels were equivalent between mutant and WT cells, and all the mutant cells expressed similar levels of SF3B1 K700E, as determined by WB analysis with a K700E-specific antibody (Fig. S1B). RNA-seq analysis of mutant cells revealed usage of cryptic 3’ splice sites (3′ss) that were approximately 10-25 nt upstream of the canonical 3′ss (Fig. S1C, S1D). This is consistent with our previous analysis of mutant SF3B1 MDS patient cells^13^ and with previous observations in other cancers^13-16^. Compared to the mouse models, 71% of the unique cryptic 3’ss in the mutant SF3B1 K562 cells overlapped with those in mutant SF3B1 MDS cells (Fig. 1B, Tables S1and S2). In addition, nine out of the top ten enriched pathways, derived from the cryptic 3’ss gene transcript list of the mutant K562 cells, overlapped with the top ten enriched pathways from the cryptic 3’ss gene transcript list of MDS patient cells (Fig. S1E). These results suggest that the mutant K562 CRISPR cells faithfully model mutant SF3B1 MDS disease.

**Figure 1.**
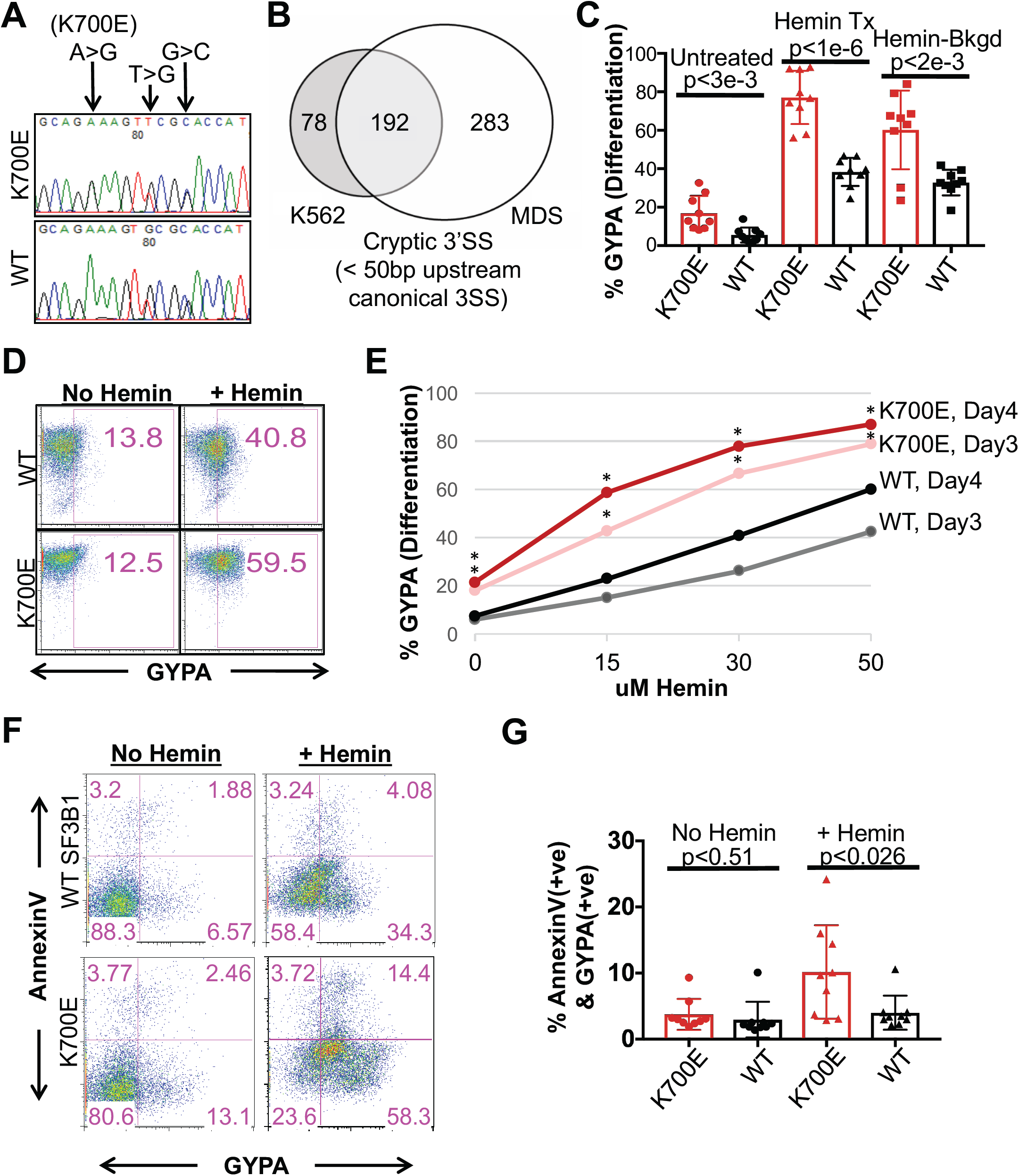
K562/SF3B1 K700E cells exhibit accelerated differentiation and increased erythroid cell death as compared to WT cells. (A) DNA chromatograms of representative K562/SF3B1 mutant and WT clones, showing K700E and two silent mutations in the *SF3B1* gene introduced into K562 cells by CRISPR. (B) Venn diagram showing the overlap of cryptic 3’ splice sites (3’ss, cut-off at q-value < 0.05, distance < 50nt to the associated canonical 3’ss) that were utilized significantly more in *SF3B1* mutants than in *SF3B1* wild-type K562 cells and MDS patients. (C) Bar graph quantifying the percentage of surface glycophorin A (GYPA) positive cells as analyzed by flow cytometry (FACS) with K700E and WT K562 cells that were treated or not with the erythroid inducer hemin (50 uM) for 3 days. Data shown represent n=6 biologically independent experiments. Each dot represents an independently derived cell clone. p-values obtained from t-tests are shown. (D) FACS plots of representative mutant and WT clones from (C) with similar background, displaying percent GYPA positive cells following hemin treatment as indicated. GYPA positivity was gated based on unstained WT and Mutant cells. (E) Line graph displaying percent GYPA positive cells vs. various concentrations (uM) of hemin for mutant and WT clones as measured by FACS after 3 and 4 days of hemin or no treatment. *, p < 0.05. (G) Bar graph specifying the percentages of Annexin V vs. GYPA expressed on the surface of K700E and WT cells as measured by FACS after hemin (50uM) or no treatment for 4 days. Representative data from n=4 independent experiments. p-values from t-tests are shown. (F) FACS plots (lower left quadrants of each plot: GYPA-AnnexinV-as undifferentiated cells; lower right quadrants: GYPA+AnnexinV-as non-apoptotic erythroids; upper right quadrants: GYPA+AnnexinV+ as apoptotic erythroids; upper left quadrants: GYPA-AnnexinV+ as apoptotic cells) of representative mutant and WT clones from (G), showing the percentages of Annexin V vs. GYPA expression under 4 days of hemin or no treatment.

We next examined whether the K700E mutation affected growth or differentiation properties of the mutant K562 cells. We observed that while the mutant cells displayed a small, unexpected growth defect compared to WT cells (Fig. S1F), there was no statistically significant difference in apoptotic cell death under normal growth conditions between K700E and WT cells (Fig. S1G). As SF3B1 mutations are associated with red blood cell dysfunction, we examined the ability of the mutant and WT K562 cells to differentiate along the erythroid lineage by treating the cells with hemin (ferriprotoporphyrin IX), an erythroid differentiation inducer known to be effective with K562 cells^33,35^. Surprisingly, the mutant cells differentiated more to erythroid cells than did the WT cells, as quantified by flow cytometric (FACS) analysis measuring glycophorin A (GYPA), a well-characterized surface marker of erythroid differentiation (Fig. 1C, 1D). While it has been reported that normal K562 cells undergo low levels of spontaneous erythroid differentiation^36^, the mutant clones had on average higher GYPA levels (Fig. 1C), indicative of an increased frequency of spontaneous differentiation. The higher percentage of overall hemin-induced erythroid differentiation, however, was not due to the increased occurrence of spontaneous differentiation, as some mutant clones had similar GYPA^+^ background as WT but had higher differentiation capacity (Fig. 1D). In fact, mutant cells differentiated more rapidly and completely to erythroid cells than did WT clones (Fig. 1E and Fig. S1H).

We next investigated the fate of the mutant K562 cells upon differentiation. Remarkably, following differentiation, the mutant cells underwent apoptosis as measured by the apoptotic surface marker AnnexinV, as the percentage (10.2 ± 6.7 in K700E vs 4.0±2.4 in WT) of apoptotic erythroid GYPA^+^ AnnexinV^+^ cells was statistically, significantly increased compared to the WT cells (Fig. 1F, 1G). Hence, our K562/SF3B1 cells recapitulate two key features of MDS cells: aberrant erythroid differentiation and increased erythroid cell death.

### p38 MAPK is specifically deregulated and only in mutant SF3B1 cells

We next wished to investigate pathways that may be of pathophysiologic significance to MDS disease. We hypothesized that a signaling pathway such as the p38 MAPK (p38) pathway, which is known to be involved in governing hematopoietic cell differentiation and apoptosis^30,36-38^, including erythropoiesis^30,32,37^, might be disrupted. To test this, we examined our panel of WT and K700E cells by WB analysis using antibodies against both p38 itself and the activated form, phospho-p38 (p-p38). Strikingly, p-p38 levels were significantly reduced in K700E cells compared to WT cells, although total p38 levels were similar (Fig. 2A; quantitation on right). Notably, of the three major MAP kinase pathways, only p38 was deactivated in K700E cells: again, using appropriate phospho-specific antibodies, we observed no differences in levels of activated JNK or ERK between mutant and WT cells (Fig. 2B, 2C).

**Figure 2.**
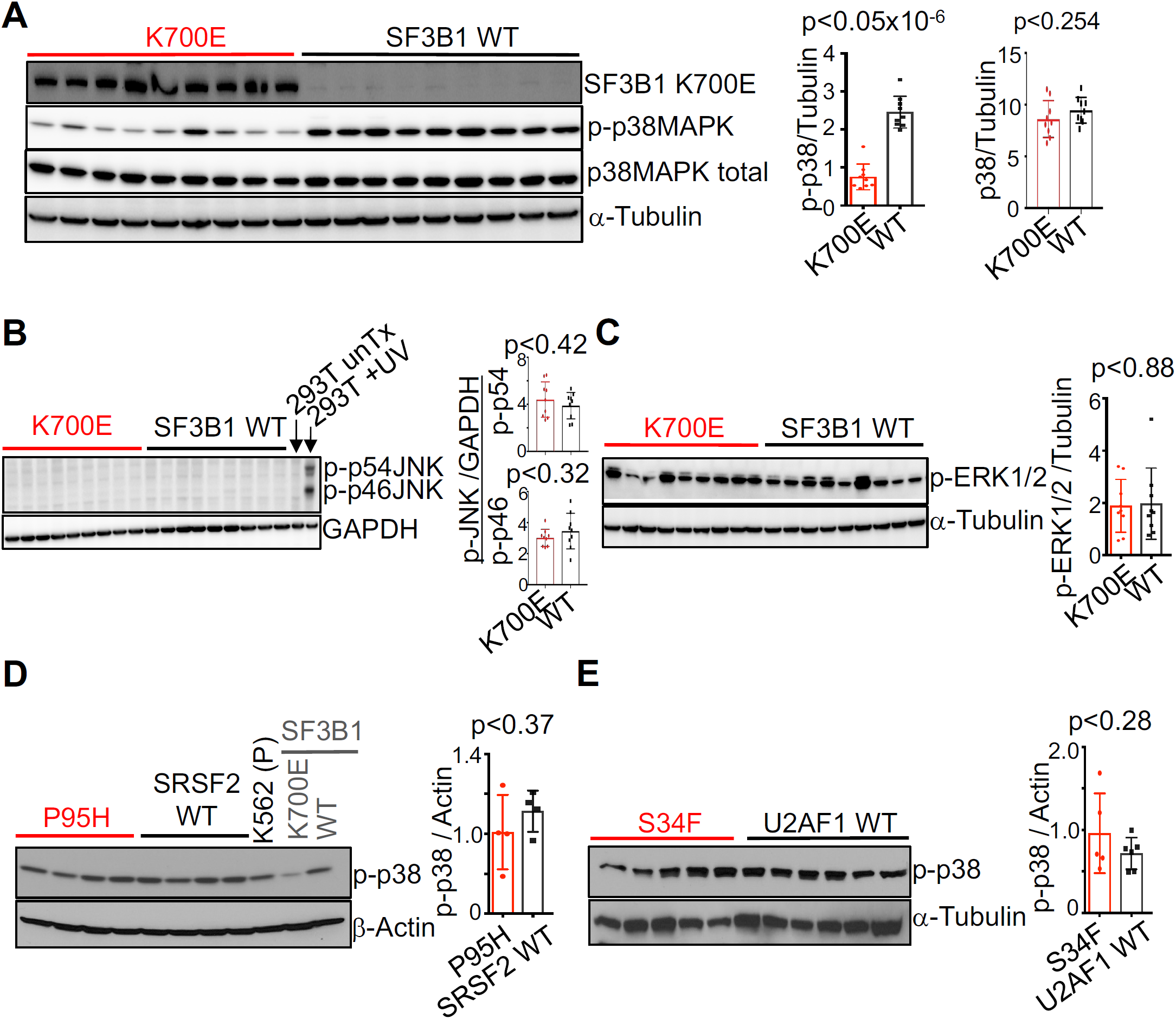
p38 MAPK is specifically deregulated and only in mutant SF3B1 cells. (A) Representative western blot (WB) images showing expression of total p38 MAPK, phospho-p38 MAPK (p-p38MAPK; p-p38), SF3B1 K700E and α-Tubulin in the nine independent mutant and nine independent WT K562/SF3B1 clones. n=5 independent experiments. Representative WB analysis of phospho-JNK (p-p46 and p-p54) and GAPDH (B), and phospho-ERK1/2 (p-ERK12) and α-Tubulin (C) in K562/SF3B1 K700E and WT clones. Representative WB analysis of p-p38 in (D) K562/SRSF2 mutant (P95H) and WT and in (E) K562/U2AF1 mutant (S34F) and WT independent clones. For (B) and (C), n ≥ 3, and for (D) and (E), n=2 independent experiments. Bar graphs are shown next to all WB images, displaying results of ImageJ-quantified, loading control-normalized protein band intensities and p-values from t-tests.

RNA splicing factor mutations occur in a mutually exclusive manner in MDS^4,39^, suggesting that the mutations could act on the same pathway. After *SF3B1*, two other genes frequently mutated in MDS encoding splicing factors are *SRSF2* and *U2AF1*. We therefore examined p38 activation in panels of U2AF1 and SRSF2 K562 cells. As with the SF3B1 cells, we used CRISPR to introduce appropriate hotspot mutations (Fig. S2 and ^40^). Interestingly, WB analysis did not reveal any differences in p-p38 levels between mutant SRSF2 P95H or U2AF1 S34F cells and their WT counterpart cells (Fig. 2D, 2E), suggesting that the p38 pathway defect is specific to SF3B1 mutant cells.

### MAP3K7 transcripts are misspliced in mutant SF3B1 cells and this is responsible for p38 deactivation

We next wished to identify the defect responsible for p38 inactivation in the SF3B1 mutant cells. We hypothesized that a mutant SF3B1-induced splicing error was involved, and made several further suppositions: first, that the ratio of misspliced to correctly spliced transcript would be sufficiently high to alter protein levels; second, that the aberrantly spliced transcript encoded a kinase or phosphatase that could directly or indirectly regulate p38; and third, that the splicing error would be found in both mutant K562 cells and MDS patient cells. When we applied these criteria, we identified the MAP kinase MAP3K7 as a likely relevant target of mutant SF3B1. MAP3K7 is a known upstream regulator of p38^41,42^, and previous studies have indicated that its transcript is indeed misspliced in MDS mutant cells^13,26,28^. Analysis of RNA-seq data from our SF3B1 mutant K562 cells and from MDS patient cells revealed extensive cryptic 3’ss usage, which was not observed in the SRSF2 and U2AF1 mutant K562 cells (Fig. 3A; Fig. S3A). Interestingly, besides in MDS, MAP3K7 is also misspliced in other cancers with SF3B1 mutations (Fig. S3B, ^6,11,43^, indicating that MAP3K7 is a common target of mutant SF3B1 regardless of tissue type. Reported as a target of NMD, MAP3K7 transcript levels were decreased in mutant SF3B1 K562 and MDS cells (Fig. 3B and^28^, but as expected, not in the mutant SRSF2 and U2AF1 K562 cells (Fig. 3B). Most importantly, WB analysis revealed that MAP3K7 protein levels were sharply reduced in all nine of the SF3B1 mutant K562 clones used in our experiments (Fig. 3C).

**Figure 3.**
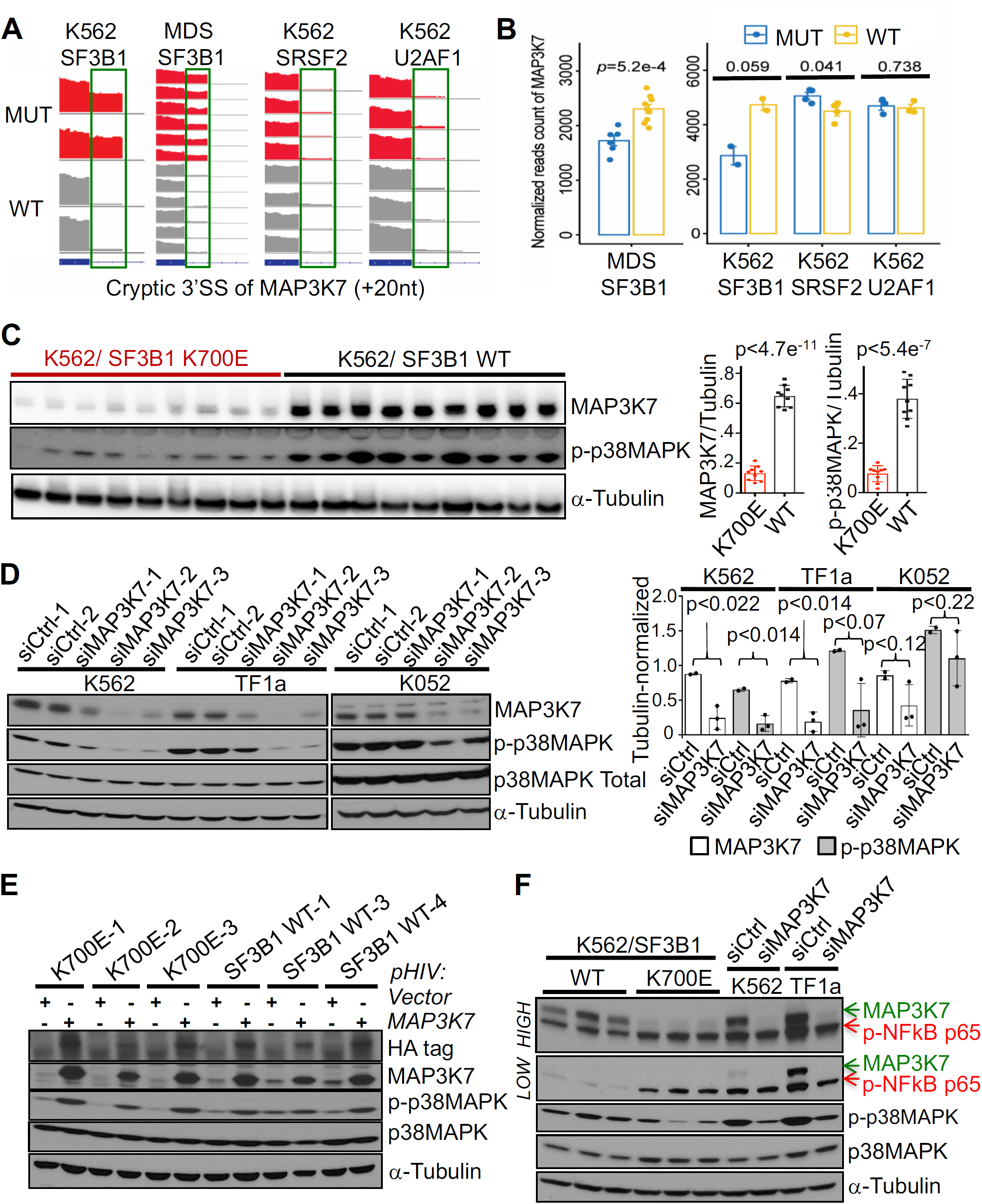
MAP3K7 transcripts are misspliced in mutant SF3B1 cells, and this is responsible for p38 deactivation. (A) RNA-seq coverage plots of *MAP3K7* cryptic 3’ss transcripts in WT and mutated samples from (left to right) K562/SF3B1, MDS patients, K562/SRSF2, and K562/U2AF1 cells. (B) Comparison of total *MAP3K7* mRNA levels (normalized reads counts) in WT and mutated RNA-seq samples from (left to right) MDS patients, K562/SF3B1, K562/SRSF2, and K562/U2AF1 cells. p-values from t-tests are shown. (C) Representative western blot images showing expression of MAP3K7, p-p38MAPK, and α-Tubulin proteins in the nine independent mutant and nine independent WT K562/SF3B1 clones. n≥6 independent experiments. Right panels, bar graphs displaying the results of ImageJ-quantified, α-Tubulin-normalized band intensities and p-values from t-tests. (D) Representative western blot analysis of the effects on p-p38 MAPK and total p38 MAPK expression in MAP3K7-KDed K562, TF1a, and K052 cells using two different negative control siRNAs (siCtrl-1/2) and three different siRNAs specific for *MAP3K7*. n≥3 independent experiments. Right panels, bar graphs displaying the results of ImageJ-quantified, α-Tubulin-normalized band intensities and p-values from t-tests. (E) Representative western blot analysis of the effects on p-p38 MAPK and total p38 MAPK expression in three WT and three mutant K562 clones expressing HA-tagged MAP3K7. n≥3 independent experiments. (F) Representative western blot analysis of the effects on phospho-NFkB p65 (p-NFkB p65), p-p38 MAPK and total p38 MAPK expression in MAP3K7-KDed K562 and TF1a cells using negative control siRNA #1 and siMAP3K7 #2 (see D). Expression of p-NFkB p65 in three mutant and three WT K562/SF3B1 is also shown. “Low” and “High” represent different exposures of the same gel. n=3 independent experiments.

We next set out to determine whether the reduced MAP3K7 levels were in fact responsible for the lower p38 activation. To this end, we knocked down (KD) MAP3K7 using three different siRNAs in parental K562 cells and in two other leukemic cell lines, TF1a and K052, and measured p-p38 levels by WB analysis (Fig. 3D). We observed significant reductions in p-p38, but not p38, levels induced by all three siRNAs. Importantly, when we reintroduced MAP3K7 into SF3B1 mutant cells using lentivirus, p-p38 abundance was restored and equivalent to that observed in similarly treated WT cells (Fig. 3E).

In addition to p38, previous studies showed that MAP3K7 can also positively regulate NF-kappaB (NFkB) under certain circumstances^44^. However, KD of MAP3K7 by siRNAs (Fig. 3F) or shRNAs (Fig. S3C) did not alter levels of phospho-NFkBp65 in parental K562 or TF1a cells, suggesting that under normal growth conditions, MAP3K7 does not regulate NFkBp65 activation. Interestingly, under the same normal growth condition, we observed that SF3B1 mutant K562 cells displayed modestly hyperactivated NFkB (Fig. 3F), consistent with results from a previous study^28^ and indicating another altered pathway of possible pathological significance in MDS. However, in contrast to the conclusions of that study^28^our data suggest that reduced MAP3K7 levels were not responsible for the NFkB activation under normal growth conditions that we observed, suggesting that another missplicing event likely underlies this response.

### Knockdown of MAP3K7 in parental K562 and normal human CD34+ cells causes increased erythroid differentiation and cell death

We next wished to determine whether reduced levels of MAP3K7 are sufficient to induce the aberrant erythrocyte differentiation and apoptosis observed in the SF3B1 mutant K562 cells. KD of MAP3K7 in parental K562 cells had no effects on cell proliferation or, similar to the mutant cells, apoptosis under normal growth conditions (Fig. S4A, 4B). However, KD of MAP3K7, by shRNA, during hemin-induced erythroid differentiation in these cells led to an increase in differentiation, as shown by an elevated percentage of total GYPA^+^ expressing cells and a subsequent reduction in undifferentiated GYPA^-^AnnexinV^-^ double negative cells (Fig. 4A). In addition, MAP3K7 KD also induced erythroid cell death (Fig. 4A), as similarly observed in the SF3B1 mutant cells.

**Figure 4.**
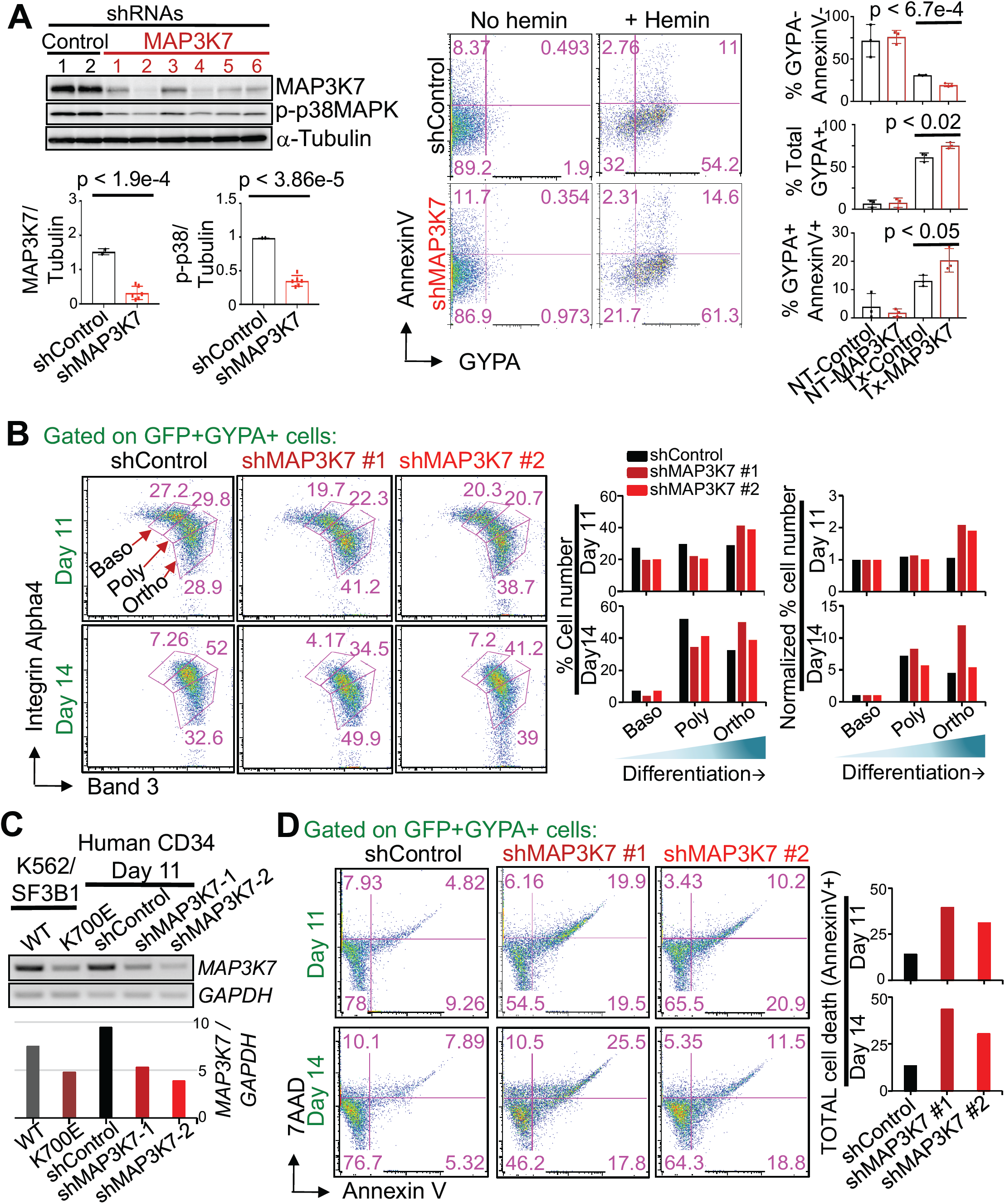
Knockdown of MAP3K7 in parental K562 and normal human CD34+ cells causes increased erythroid differentiation and cell death. (A) Images (from left to right) of representative western blot analysis of GFP-sorted, shRNA-mediated MAP3K7 KD (two different negative control and six different MAP3K7 shRNAs) in parental K562 cells, representative two-color FACS plot of AnnexinV vs. GYPA from a control and shMAP3K7, and bar graphs showing the averages and standard deviations of (from top to bottom) % GYPA-Annexin-(undifferentiated cells), % total GYPA+(non-apoptotic erythroids), and % GYPA+AnnexinV+ cells (apoptotic erythroids) after three days of treatment with 50 uM hemin from three independent experiments. Bar graphs displaying the results of ImageJ-quantified, α-Tubulin-normalized MAP3K7 and p-p38 band intensities in western blot analysis are shown below the WB. p-values from t-tests are labelled on all bar graphs. (B) Representative FACS plots showing surface expression of Integrin Alpha-4 and Band 3 from day 11 and day 14 shRNA-mediated *MAP3K7* KD (one negative control and two different MAP3K7 shRNAs) in human erythropoietin (EPO)-induced CD34+ cells that were CD45- and double positive GFP+GYPA+ (= erythroblasts cells). Erythroblast stages are depicted and labelled (Baso, basophilic normoblast; poly, polychromatophilic normoblast; ortho, othrochromatic normoblast). Right panels, bar graphs, quantifying the percentage of erythroblasts in each stage, are shown as well as Baso-normalized percent erythroid cells for comparison. n=2 independent experiments. (C) RT-PCR showing day 11 shRNA-mediated *MAP3K7*-KDed expression in day 11 EPO-induced CD34+ cells. Abundance of *GAPDH*-normalized *MAP3K7* expression is depicted in the bar graph below. (D) Representative FACS analysis of cell death via 7AAD and Annexin V from (C) day 11 and day 14 *MAP3K7* KDed, human EPO-induced, CD45-GFP+GYPA+ erythroblasts. Bar graphs, specifying the percentage of total erythroblast cell death, are shown. n=2 independent experiments.

Because all the functional studies described above were done with K562 erythroleukemia cells, we next wanted to evaluate the effects of MAP3K7 KD on erythroid differentiation in normal, nontransformed cells. For this, we used human CD34^+^ adult blood stem cells, induced the cells to differentiate along the erythroid lineage with erythropoietin (Epo), and then employed a refined, three-erythroid surface marker (GYPA, Integrin Alpha4, Band 3) method that allows for determination of the four stages of erythroblast maturation: proerythroblast, basophilic normoblast, polychromatophilic normoblast, and orthochromatic normoblast^45^. Elucidating these four stages was done by FACS analysis by first gating on the GYPA marker that is present on cells during all four erythroblast stages and then examining for surface levels of Integrin Alpha-4 and Band 3; expression of Integrin Alpha-4 decreases with development while surface expression of Band 3 increases with maturation^45^. Importantly, MAP3K7 KD, using two different shRNAs, again resulted in accelerated differentiation: Fewer KD CD34+ cells were present at early staged erythroblasts and more at late stages compared to the negative control shRNA (Fig. 4B, 4C). In addition, we observed more erythroid cell death in the two MAP3K7 KD cells than in the negative control cells (Fig. 4D).

The above experiments together provide strong support for our hypothesis that the decreased MAP3K7 abundance in K700E cells was responsible for the accelerated erythroid development and subsequent cell death. This is consistent with previous reports documenting erythroid expansion in the bone marrows of RARS patients^7,19,46^, in which, as mentioned earlier, *SF3B1* is mutated in 65-83% of this MDS subtype. In fact, it was reported that MDS patients harboring *SF3B1* mutations have increased erythroid activity and accumulation of erythroblasts in their bone marrows^8^.

### Premature downregulation of GATA1 in differentiated K700E or MAP3K7 KD cells underlies the accelerated differentiation and erythroid cell death

We next investigated the mechanism by which reduced levels of MAP3K7 caused accelerated erythroid differentiation followed by apoptosis. We examined the expression levels of some major erythroid transcription factors by WB (Fig. 5A) or qPCR (Fig. S5) in mutant and WT cells during erythroid differentiation. Based on the expression patterns of these transcription factors and their connection or lack thereof to p38, we hypothesized that the transcription factor GATA1, known to be a master regulator of erythroid differentiation and a downstream target of p38 ^47-49^, is misregulated in mutant SF3B1 cells. To investigate this, we first examined GATA1 expression during a time course of hemin-induced erythroid differentiation by WB analysis, using two representative K700E and WT K562 clones (Fig. 5A). The two WT clones showed that GATA1 expression decreased during differentiation (Fig. 5A), which is consistent with previous studies showing that during erythroblast differentiation, GATA1 expression is required to be downregulated for erythroid maturation ^50^. Interestingly, GATA1 expression in the two K700E clones was also downregulated, but to a greater extent and more rapidly than in the two WT clones (Fig. 5A). To confirm these results, we examined our full panel of nine K700E and nine WT clones after three days of differentiation, and observed that GATA1 was indeed more downregulated in K700E than in WT cells (Fig. 5B). To show that the reduced MAP3K7 levels in the mutant K562 cells were responsible for the greater downregulation of GATA1 during differentiation, we examined GATA1 levels in MAP3K7 KD parental K562 cells, and again observed a greater decrease in GATA1 expression than in the negative control cells (Fig. 5C).

**Figure 5.**
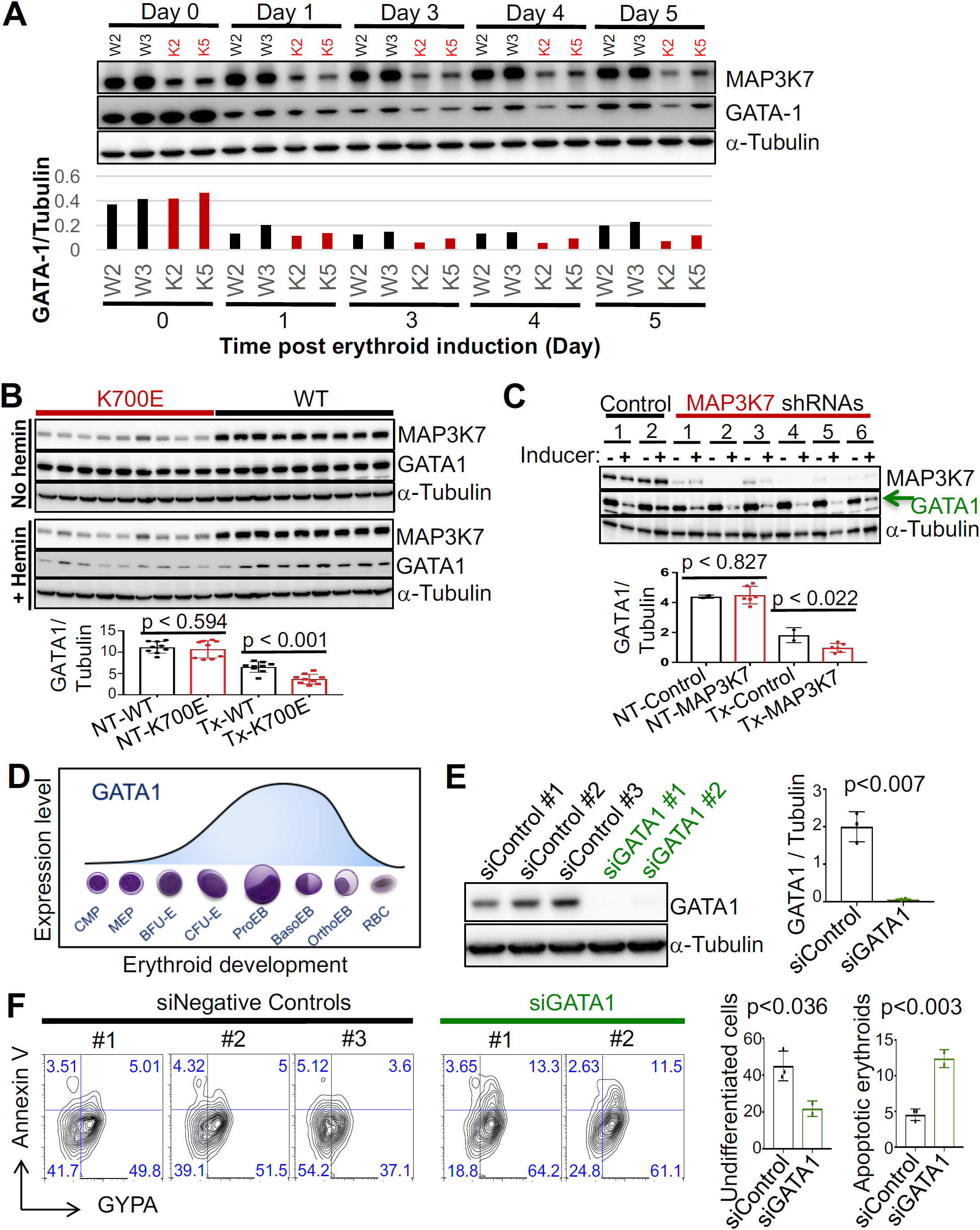
Premature downregulation of GATA1 in differentiated K700E or MAP3K7 KD cells underlies the accelerated differentiation and erythroid cell death. (A) Western blot analysis (top) and quantifying bar graph (bottom) showing expression of GATA-1 during a 5-day time course of treatment with 50 uM hemin to induce erythroid differentiation in two representative K700E (K2 and K5) and two representative WT (W2 and W3) K562/SF3B1 clones. (B) Representative western blot images showing GATA-1 protein expression in the nine mutant and nine WT K562/SF3B1 clones that were treated or not with 50 uM hemin for three days. Bar graph displaying the results of ImageJ-quantified, α-Tubulin-normalized GATA-1 band intensity and p-values from t-tests. n=4 independent experiments. (C) Representative western blot image of GATA-1 expression in shRNA-mediated MAP3K7 KDed parental K562 cells that were treated or not with 50 uM hemin for three days. n=3 independent experiments. (D) Illustration (adapted from Moriguchi and Yamamoto, 2014) showing GATA1 expression during the course of erythroid development from myeloid progenitor cell (CMP) to mature red blood cell (RBC). MEP, megakaryocyte-erythroid progenitor; BFU-E, Burst-forming unit-erythroid; CFU-E, colony-forming unit-erythroid; ProEB, Proerythroblast; BasoEB, basophilic erythroblast/normoblast; OrthoEB, orthochromatic erythroblast/normoblast. (E) Western blot image showing GATA-1 KDed (two different GATA-1 and three different negative control siRNAs) expression in parental K562 cells and (F) its effects on erythroid differentiation and erythroid apoptosis via FACS analysis of Annexin V vs. GYPA after 3 day of 50 uM hemin treatment. Bar graphs (right) indicate % undifferentiated cells (double negative GYPA-AnnexinV-) and % apoptotic erythroid cells (double positive GYPA+AnnexinV+). p-values from t-tests are shown. n=3 independent experiments. GYPA positivity was gated based on unstained parental K562 cells.

During erythroid differentiation, the requirement for GATA1 expression is biphasic: there is a steadily increasing demand for GATA1 by erythroid-committed progenitor cells that peaks prior to the onset of the erythroblast stages; during the erythroblast stages, GATA1 expression steadily decreases to enable erythroblasts to properly mature (Fig. 5D, adapted from Moriguchi and Yamamoto, 2014). Based on our findings, we hypothesized that GATA1 expression in mutant cells decreases faster and more extensively than in WT cells during the erythroblast stages, thus causing the accelerated differentiation and subsequent apoptosis. To test this hypothesis, we pre-treated parental K562 cells with hemin to initiate early erythroid differentiation, and then KDed GATA1 in the hemin pre-treated cells. Strikingly, GATA1 KD after hemin pretreatment resulted in increased erythroid differentiation and subsequent apoptotic cell death compared to the negative control cells (Fig. 5E, 5F), completely analogous to what we observed in SF3B1 mutant cells or following MAP3K7 KD. Together, our data strongly supports the hypothesis that the earlier and more extensive downregulation of GATA1 in differentiated K700E cells underlies the accelerated differentiation and erythroid cell death we observed.

### Mutant SF3B1 MDS patient cells display lower levels of phospho-p38 and MAP3K7, accelerated differentiation, and increased erythroid cell death compared to patients with WT SF3B1

Our experiments have shown that the aberrant differentiation followed by apoptosis characteristic of MDS can be recapitulated in SF3B1 mutant K562 cells. However, we next wished to determine whether SF3B1 MDS involves the dysregulation of the MAP3K7-p38 pathway we have described in this report. To investigate this, we first examined expression of p-p38 and MAP3K7 in SF3B1 MDS bone marrow mononuclear cells (BMMNCs) by WB analysis (Fig. 6A). Consistent with the SF3B1 mutant K562 cells, p-p38 and MAP3K7 levels were significantly reduced in mutant SF3B1 MDS BMMNCs compared to WT SF3B1 MDS patient BMMNCs (Fig. 6A). To ascertain the erythroid differentiation and cell death properties of the mutant SF3B1 MDS cells, we examined the erythroblast stages of mutant SF3B1 MDS and normal bone marrow cells using the three-erythroid surface marker method described above ^45^. Similar to what we observed with MAP3K7-KD CD34^+^ cells, mutant SF3B1 MDS patient bone marrows had less early staged erythroblasts and more late staged erythroid cells compared to the normal healthy bone marrow control cells (Fig. 6B). This is consistent with Figure S6A, which was generated by replotting only the normal controls and MDS SF3B1 K700E patient data reported by ^51^. The data in Figure S6A also revealed less early staged erythroblasts and more late staged erythroid cells in the SF3B1 mutant samples. Additionally, when we measured erythroid cell death by Annexin V staining, the mutant SF3B1 MDS GYPA+ cells showed more total cell death than did those from normal bone marrow (Fig. 6C). Together, our data explains the anemic phenotype of mutant SF3B1 MDS patients and demonstrates that the MAP3K7-p38 pathway we elucidated using mutant SF3B1 K562 cells explains the phenotypes of mutant SF3B1 MDS patient bone marrow cells.

**Figure 6.**
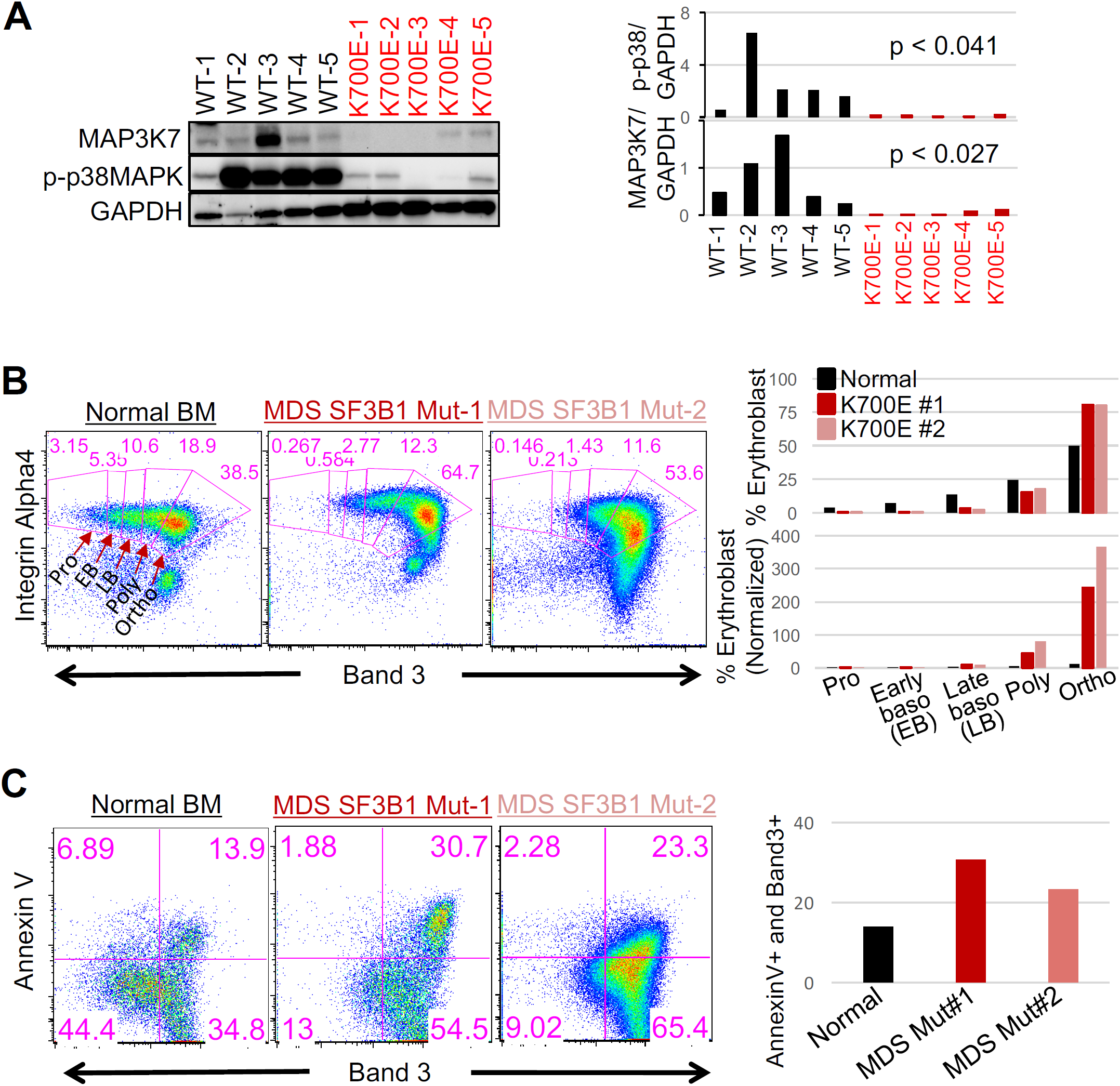
Mutant SF3B1 MDS patient cells display lower levels of phopho-p38 and MAP3K7, accelerated differentiation, and increased erythroid cell death compared to patients with WT SF3B1. (A) Western blot analysis showing MAP3K7 and p-p38 MAPK expression in MDS bone marrow (BM) mononuclear cells from five patients with WT and five patients with K700E SF3B1 mutation. Right panel, bar graphs quantifying the results of WBs and p-values from t-tests are shown. (B) Representative FACS plots showing the erythroblast profiles of primary BM cells from SF3B1 K700E MDS patients and normal healthy individual. The isolated CD45-BM cells were stained with 3 erythroid markers: GYPA, Integrin Alpha-4, and Band3. FACS plot of Integrin Alpha-4 vs. Band 3 on GYPA+ BM cells. Erythroblast stages are depicted and labelled. Bar graphs, quantifying the percentage of nucleated erythroblasts in each stage as a total of 100%, are shown as well as Proerythroblast (Pro)-normalized percent erythroid cells for comparison. Three SF3B1 K700E MDS patients and three normal healthy individuals were profiled. Pro, proethroblasts; EB, early basophilic erythroblasts; LB, late basophilic erythroblasts; Poly, polychromatic erythroblasts; and Ortho, orthochromatic erythroblasts. (C) Representative FACS analysis of erythroblast cell death via Annexin V vs. Band 3 from (B). Bar graph quantifies the percentage of late-stage erythroblast cell death (AnnexinV+Band3+).

### Other *SF3B1* mutations lead to extensive MAP3K7 missplicing

Our studies focused on the most common hotspot mutation found in MDS patients, K700E. Since we demonstrated that reduced levels of MAP3K7 alone, in K562 and normal CD34+ cells, was sufficient to cause enhanced erythroid development, expansion and cell death, any SF3B1 mutation that induces missplicing of MAP3K7 almost certainly behaves identically to K700E. Analysis of our own and other RNA-seq datasets showed that all other MDS *SF3B1* mutations that we examined induced extensive missplicing of MAP3K7, ranging from 50-90% (Fig. S6B and ^18,52^. This includes four additional hot spot mutations (E622, R625, H662, and K666) and one non-hotspot mutation (D781G). Thus, the decreased p38 activation and the resultant phenotypes reported here very likely apply to all or virtually all MDS mutant SF3B1 patients. In support of our hypothesis that other non-K700E SF3B1 mutations behave identically to K700E in terms of greater erythroid development and expansion, we again reanalyzed the data reported by Ali et al. (2018) and replotted only normal controls and MDS samples with non-K700E SF3B1 mutations (3 E622, 4 R625, 4 H662, 2 K666, and 1 D781G) (Fig. S6C). The resultant plot showed less early staged but more late staged erythroblasts in the non-K700E SF3B1 mutant cells, which is consistent with our data (see Fig. 6B and Fig. S6A).

## DISCUSSION

*SF3B1* is the most frequently mutated gene in MDS^9^, and such mutations are clinically associated with a distinct type of anemia that is characteristically of the RARS subtype. However, the underlying molecular mechanism responsible for anemia has been unknown. In the present study, we found that SF3B1 K700E-induced missplicing of MAP3K7 mRNA is responsible for p38 deactivation and ultimately for faster and greater GATA1 down-regulation. This in turn causes aberrant differentiation and apoptotic erythroid cell death, as found in both our mutant SF3B1 K562 cells and MDS patient cells. Our findings provide a detailed mechanism explaining the origins of anemia in MDS patients harboring *SF3B1* mutations (see model in Fig. 7D). Below we discuss additional implications of our results with respect to MDS, more details concerning the p38-based pathway we discovered, and how our findings may have therapeutic implications.

**Figure 7.**
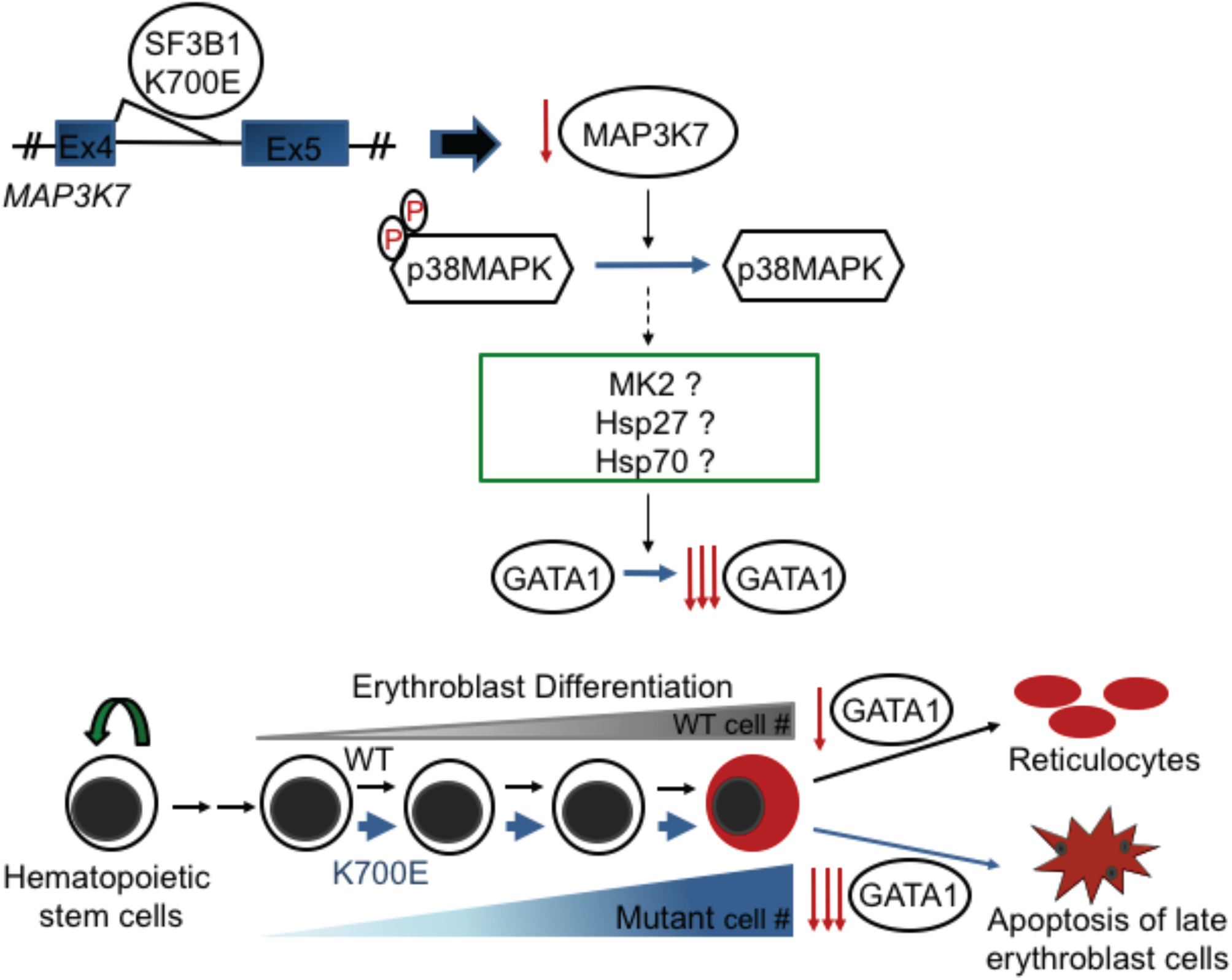
A model for how *SF3B1* mutations cause anemia. SF3B1 K700E, and other hotspot mutations, induce an error in splicing of MAP3K7 transcripts, resulting in reduced levels of MAP3K7. This leads to a reduction of inactivated, phosphorylated p38MAPK (p-p38), which in turn affects downstream potential targets such as MAPKAPK2 (MK2), HSP27, and/or HSP70. Ultimately, this causes faster and greater downregulation of GATA1, resulting in accelerated differentiation and erythroid cell death, thereby explaining the anemia that characterizes MDS patients harboring SF3B1 mutations.

The presence of ringed sideroblasts (≥ 15%) in MDS is clinically associated with BM hypercellularity, another hallmark feature of RARS MDS^53^. Our findings explain how this hypercellular phenotype arises in the BM of mutant SF3B1 MDS patients through accumulation of erythroblasts. During erythroid development from progenitor cells to late staged erythroblasts, there is a successive expansion of these nucleated cells as they mature^54^. Thus, late staged erythroblasts are the most numerous nucleated erythroid cells in normal bone marrow, as also shown in our differentiated CD34+ cells and normal bone marrow. However, MAP3K7-KD CD34+ cells and mutant SF3B1 bone marrow had an even higher abundance of late staged erythroblasts, indicating greater erythroid development and activity. Overall, our findings are in agreement with, and provide a mechanistic explanation for, reports showing aberrant erythroid maturation, accumulation of erythroid precursors/BM hypercellularity, and more erythroid cell death in the bone marrow of RARS MDS patients^20,21,53^. In fact, it has been shown that there are significantly more erythroblasts in the bone marrows of MDS patients with mutant vs WT SF3B1^8^ and that the *SF3B1* mutant allele burden positively correlates with the percentage of BM erythroblasts^7^.

The expression profile of GATA1 during the development of progenitor cells to reticulocytes indicates that GATA1 expression must decrease for erythroblasts to mature(Fig. 5D, an adaptation^50^). Thus, our observation that aberrantly faster and greater downregulation of GATA1 occurs in mutant SF3B1 cells provides an explanation for the accelerated differentiation phenotype in the mutant SF3B1 K562 cells and, more importantly, MDS patient cells. Consistent with this notion, it had been demonstrated that both GATA1 mRNA and protein levels were reduced in purified day 7 and day 14-cultured GYPA+ erythroid RARS cells as compared to those in normal bone marrow cells^55^. Furthermore, decreased GATA1 expression in erythroblasts has been linked to ineffective hematopoiesis^56^.

In normal erythroid development, terminal differentiation is accompanied by cessation of cell proliferation^57^. However, we found that the cell proliferation program, as demonstrated by the proliferation marker Ki67, was more active in mutant SF3B1 K562 cells than in WT cells (Fig. S7A). Consistent with this, levels of soluble transferrin receptor, which is related to the cellular iron uptake and erythroid proliferation rate^58^, have been reported to be elevated in SF3B1 mutant MDS patients^7,8^. These findings, in conjunction with increased abundance of late staged erythroblasts in mutant SF3B1 MDS and MAP3K7-KD cells, provide a functional explanation for the accumulation of erythroblasts that occurs in patients with RARS MDS with *SF3B1* mutations. The basis for the uncoupling of terminal differentiation from cell-cycle arrest could be a result of the faster and greater downregulation of GATA1. During erythroid differentiation, GATA-1 has been shown to directly activate expression of p21^59^, a known negative regulator of the cell cycle. As GATA1 was reduced faster and to a greater extent in mutant SF3B1 and MAP3K7-KD cells, leading to accelerated differentiation, p21 is likely not activated sufficiently to turn off cell proliferation in these cells. The conflicting signals of proliferation and terminal differentiation, where cell-cycle arrest must occur, could trigger apoptosis^60^. Alternatively, it has been shown that GATA-1 can stimulate the expression of *Bcl-xL*^61-63^, which encodes a pro-survival Bcl-2 family protein and is known to be upregulated in erythroblasts^64,65^. The faster and greater downregulation of GATA-1 thus might not activate *Bcl-xL* sufficiently to confer erythroblast survival, providing another explanation for the apoptotic erythroid phenotypes we observed in mutant SF3B1 MDS cells.

Another question concerns the mechanism leading from reduced p-p38 levels to the earlier and greater GATA1 downregulation during differentiation in mutant SF3B1 cells. It has been shown that Caspase 3-mediated cleavage of GATA1 results in GATA1 downregulation, and this is important for normal erythroid maturation^48^. However, Caspase 3 cleavage of GATA1 is normally blocked in the early stages of differentiation by association of GATA1 with Hsp70 in the nucleus, allowing for GATA1 activation of target genes^48^. Interestingly, Hsp70 nuclear localization has been shown to be defective in MDS, leading to premature GATA1 cleavage^66^. Although the basis for the aberrant localization of Hsp70 is not known, Hsp70 can directly interact with p38 and can also influence the phosphorylation of MAPKAPK2 (MK2), a downstream target of p38^67^ and whose expression is downregulated in mutant K562 cells (Fig. S7B). Thus, the reduced p38 activation we have characterized may underlie this defect. Another possible intermediate effector between p38 and GATA1 is Hsp27, which has been shown to be a downstream target of p38 and can regulate GATA1 protein abundance^47^. Interestingly, Hsp27 is also downregulated in mutant K562 cells (Fig. S7). However, future experiments will be required to elucidate the precise mechanism.

A number of previous studies designed to understand the functional consequences of *SF3B1* mutation have used knock-in K700E mouse models. Such studies demonstrated that the presence of dysplastic erythroid cells is a shared phenotype between SF3B1 mutant MDS patient samples and mutant mice^17,18^. However, these mice were not able to recapitulate certain other hallmark features of mutant SF3B1 MDS, or specifically RARS MDS, such as accumulation of erythroblasts, apoptotic erythroid progenitors and erythroblasts, and ringed sideroblasts. This inability to recapitulate many of the key features of MDS, particularly RARS, could be attributed to the minimal overlap between aberrantly spliced transcripts in MDS patients and mice (~5-10%^7,18^). Thus, the differences between mouse and human support the idea that the principal features of RARS MDS are caused by mutant SF3B1-induced missplicing.

In addition to qualitative differences in missplicing between mice and humans, there are also quantitative difference in the extent of missplicing of specific transcripts. For example, Lee et al. (2018) found more overlap (21%) in missplicing between SF3B1 mutant MDS patients and K700E knock-in mice. Importantly, one of the overlapped transcripts was *MAP3K7*. However, Lee et al. (2018) found that only ~5-10% of *MAP3K7* transcripts were misspliced in the mutant mice, while we observed 51-72% *MAP3K7* missplicing in both our SF3B1 K700E K562 cells and in MDS patient cells. While protein levels were not measured, the low abundance of misspliced *MAP3K7* mRNA in the mutant mice was most likely not sufficient to affect protein levels sufficiently to account for any erythroid phenotypes. This could be another reason for the limited overlap in erythroid phenotypes between K700E knock-in mice and SF3B1 mutant MDS patients.

*SF3B1* mutations are frequent in MDS, but *MAP3K7* mutations in MDS have not been described. This is especially notable because *MAP3K7* mutations and gene aberrations occur in other cancers^68,69^. However, as we have shown here, reduced levels of MAP3K7, resulting from either missplicing or KD, had an anti-cancer, tumor suppressor-like effect during erythroid differentiation by causing cell death as an endpoint. This may explain the lack of *MAP3K7* mutations in MDS, which often transforms to AML. MAP3K7 deficiencies in other cancers must affect different pathways than we have shown here for MDS.

The above begs the question, how do *SF3B1* mutations lead to leukemogenesis in MDS? While *SF3B1* mutations that induce MAP3K7 missplicing had deleterious effects on differentiated erythroid cells, our work and that of Lee et al. (2018) showed that under normal, unstimulated growth conditions, these same mutations can also lead to activation of NFkB, which is known to promote malignancy^70^. In addition, it was shown that *SF3B1* mutations can alter splicing of two transcripts, encoding BRD9 ^71^ and DVL2^*43*^, that can contribute to tumorigenesis in uveal melanoma and chronic lymphocytic leukemia, respectively. BRD9 is a component of a BAF chromatin remodeling complex while DVL2 acts in the Notch signaling pathway. However, there is no evidence that either of these pathways are relevant to MDS. These anti- and pro-malignant effects of *SF3B1* mutations might both explain why *SF3B1* but not *MAP3K7* mutations are associated with MDS, as well as at least partially explain the generally good prognosis that characterizes patients with *SF3B1* mutations^5,6,8,9^. In addition, patients with *SF3B1* mutations have a low percentage (>5%) of bone marrow blasts/stem cells^2,8^, which most likely determines their better clinical outcomes. Perhaps related to this, we observed an increased percentage of spontaneous erythroid differentiation in our mutant SF3B1 K562 cells under normal growth conditions. Interestingly, it has been reported that some erythroid pathway-related transcripts, encoding heme biosynthetic enzymes, globins and transcription factors, are upregulated in mutant SF3B1 MDS CD34+ cells^52^. These could provide an explanation for the low percentage of bone marrow blasts seen in MDS patients carrying *SF3B1* mutations.

Our findings suggest important new avenues for developing novel therapeutic approaches for treating MDS. For example, as faster downregulation of GATA1 underlies the accelerated erythroid differentiation and apoptosis, restoring proper expression of GATA1 could provide an opportunity for therapeutic intervention. In addition to GATA1, our study provides other opportunities for therapeutic intervention. These include activation of p38 to physiological levels using small molecule activators, usage of splice site-switching antisense oligonucleotides to prevent MAP3K7 missplicing, and using small molecules to rescue the aberrant differentiation and apoptotic erythroid cells. As MDS has become the most frequent malignancy in the elderly, finding a cure or effective treatment for these disorders is urgently needed. Our findings, by providing a mechanistic explanation for the anemia that characterizes SF3B1 MDS, which affects approximately a quarter of the MDS patient population, have the potential to significantly accelerate such therapeutic efforts.

## ACKNOWLEDGEMENTS

We thank Matthew B. Thomsen and Dr. Govind Bhagat for histological advice/analysis. We also thank Dr. Jiquang Wang for his advice on computational analysis. In addition, we appreciate the technical assistance and/or figure preparation help from the following people: Cao Chen, Muhammad Mumtaz, Jacob Hess, and Lena Huang. Lastly, we thank Dr. Donald Rio of University of California, Berkeley for critical reading of the manuscript. Most of the cell cytometric sorting and analysis were done at the HICCC and C2B2 flow cytometry cores (NIH grants: P30CA013696 and S10OD020056); NIH R35 GM118136 to J.L.M.; the Edward P. Evans Foundation grant to J.L.M.; R.R. is supported by NIH grants (R01 CA185486, R01 CA179044, U54 CA193313 and U54 209997) and NSF/SU2C/V-Foundation Ideas Lab Multidisciplinary Team (PHY-1545805). Z.L. is supported by NIH grant P01CA087497; HL140625 grant to X.A.

## AUTHOR CONTRIBUTIONS

J.L.M. and S.M. provided study supervision. Y.K.L. designed the study and wet experiments. Z.L. and R.R. designed the computational experiments. Z.L. and A.P. performed computational analysis. Y.K.L. and X.W. performed the human CD34 experiments. Y.K.L. performed all the other experiments. A.M.A., A.R., J.Z. and X.A. contributed reagents. Y.K.L, Z.L., J.Z. and J.L.M prepared the manuscript.

## DECLARATION OF INTERESTS

All authors declare no conflict of interest. Y.K.L., J.Z., J.L.M. and S.M. are supported in part by a grant from Celgene Pharmaceutical Company (currently Bristol Myers Squibb); none of the work is directly related to the current manuscript.

Further information and requests for resources and reagents should be directed to and will be fulfilled by James L. Manley.

## METHODS

### Experimental model and subject details

K562, TF1a (ATCC), and K052 (DSMZ) cells were cultured in Iscove’s Modified Dulbecco’s Medium (IMDM; Gibco, cat # 12440-053) supplemented with 10% fetal bovine serum (FBS; Seradigm, cat # 89510-186) in a 37°C, 5% CO_2_ incubator. Adult human peripheral CD34+ cells were purified from donated whole blood (New York Blood Center Blood Bank). MDS bone marrow aspirates were obtained from patients, who were seen at New York Presbyterian Hospital/Columbia University Irving Medical Center and provided informed consent for the study that was approved by the institutional review board of Columbia University and was in accordance with the Declaration of Helsinki. Fresh, normal, healthy human bone marrow mononuclear cells were purchased from PPA Research Group (Cat # 15-00073). Fresh, healthy human bone marrow aspirates were purchased from Lonza (Cat # 1M-105).

### Generation of knock-in cell lines using CRISPR/Cas9 genome editing

The SF3B1 K700E mutation along with two silent mutations was knocked into K562 cells using CRISPR/Cas9 technology as previously described (Zhang et al., 2015). For true isogenic controls, only the two silent mutations were knocked into K562 cells to generate independent WT control cell lines. The human SF3B1 CRISPR guide RNA was 5’-TGGATGAGCAGCAGAAAGTTcgg-3’. The single-stranded oligodeoxynucleotides (ssODNs, Integrated DNA Technologies) for WT and K700E SF3B1were 5’-AATGTTGGGGCATAGTTAAAACCTGTGTTTGGTTTTGTAGGTCTTGTGGATGAGCAGCAGAAAGTGCGCACCATCAGTGCTTTGGCCATTGCTGCCTTGGCTGAAGCAGCAACTCCTTATGGTATCGAAT-3’ and 5’-AATGTTGGGGCATAGTTAAAACCTGTGTTT GGTTTTGTAGGTCTTGTGGATGAGCAGCAGGAAGTGCGCACCATCAGTGCTTTGGCC ATTGCTGCCTTGGCTGAAGCAGCAACTCCTTATGGTATCGAAT-3’, respectively. Mutation knock-in was confirmed using PCR amplification of genomic DNA (forward primer:5’-GTTGATATATTGAGAGAATCTGGATG-3’; and reverse primer: 5’-AAATCAAAAGGTAATTGGTGGA-3’) and DNA sequencing.

### MAP3K7 expression constructs

5’ hemagglutinin (HA) tag-containing MAP3K7 forward primer (5’-cagtGGGCCCaccATGTA CCCATACGATGTTCCAGATTACGCTAGCGGCCGCATGTCTACAGCCTCTGCCG-3’) and its reverse primer (5’-ATAggatccTCATGAAGTGCCTTGTCGTTTC-3’) were used to amplify MAP3K7 from pDONR223-MAP3K7 plasmid (Addgene plasmid #23693) and cloned into the NotI and BamHI sites of lentiviral vector pHIV-Zsgreen (Addgene plasmid #18121) to generate the pHIV-Zsgreen-5’HA tagged MAP3K7 construct. pLKO.1-eGFP was generated from pLKO.1-puro (Sigma) by replacing the coding sequence of the puromycin resistance gene with that of eGFP using restriction sites BamHI and KpnI (5’-CGCGGATCCACCGGAGCTT<u>ACCATGGTGAGCAAGGGCGA-3’; 5’-</u>GGGGCGGGCG<u>TTACTTGTACAGCTCGTCCATG-3’)</u>. Short hairpin RNAs (shRNAs) for negative controls and for targeting mRNAs of MAP3K7 were each cloned into pLKO.1-eGFP plasmid. The shRNA sequences were: shNegative Control #1, 5’-ccgg CAACAAGATGAAGAGCACCcctctcaacactggGGTGCTCTTCATCTTGTTGtttttg-3’; shNegative Control #2, 5’-ccggTTCTCCGAACGTGTCACGTcctctcaacactgg ACGTGACACGTTCGGAGAAtttttg-3’; shMAP3K7 #1, 5’-ccgg CCTGAAACCACCAAACTTAcctctcaacactggTAAGTTTGGTGGTTTCAGGtttttg-3’; shMAP3K7 #2, 5’-ccggCATGCAACCCAAAGCGCTAcctctcaacactgg TAGCGCTTTGGGTTGCATGtttttg-3’; shMAP3K7 #3, 5’-ccggGTGTTTACAGTGTTCCCAA cctctcaacactggTTGGGAACACTGTAAACACtttttg-3’; shMAP3K7 #4, 5’-ccggTGGCTTATCTTACACTGGAcctctcaacactgg TCCAGTGTAAGATAAGCCAtttttg-3’; shMAP3K7 #5, 5’-ccgg GAGGAAAGCGTTTATTGTAcctctcaacactggTACAATAAACGCTTTCCTCtttttg-3’; shMAP3K7 #6, 5’-ccggCCCAATGGCTTATCTTACAcctctcaacactgg TGTAAGATAAGCCATTGGGtttttg-3’. All PCR-amplified products were confirmed by DNA sequencing, and all plasmids were validated by digestion with multiple restriction enzymes and partial DNA sequencing.

### Viral production

For the re-expression and KD studies, lentiviral pHIV-Zsgreen and retroviral MSCV_UBC viral vectors encoding MAP3K7 and pLKO.1_eGFP lentiviral vectors containing the six MAP3K7 shRNAs or negative control shRNAs were constructed as described above. Lentiviral vectors, along with the packaging plasmid PsPAX2 (Addgene #12260) and the envelope plasmid pCMV-VSV-G (Addgene #8454), were co-transfected into HEK293T cells that were cultured in Dulbecco’s Modified Eagle’s Medium (Gibco, cat # 11965092) supplemented with 10% FBS (VWR Seradigm, cat # 89510-186) in a 37°C, 5% CO_2_ incubator. For retroviral production, retroviral vectors were co-transfected with pCGP packaging and pCMV-VSV-G (Addgene #8454) envelope plasmids into HEK293T cells. Viral supernatant was harvested at 48h and 72h to transduce cell lines and human CD34+ cells using standard protocols. Cells were sorted for GFP positivity before proceeding with the assays (see more details below).

### siRNA transfection

siRNA transfections of K562, TF1a, and K052 suspension cells were carried out using the Neon Transfection System (Thermo Fisher Sci) with the Neon 100uL tip kit that includes the R buffer for electroporation (Thermo Fisher Sci). One million cells were resuspended in 100uL of R buffer, containing 1uL of 100uM siRNAs. Transfection was performed following the manufacture’s protocol. Electroporated cells were immediately placed in 2mL complete media for 24h. After 24h, an additional 8mL of complete media was added to the 2mL cells for 24h. Cells were harvested for subsequent assays. For GATA1 KD, 17.5-18.5h hemin (50uM final)-pretreated K562 cells were used for electroporation. Transfected cells were immediately put into 2mL complete media containing 50uM hemin for 24h. After 24h, an additional 13mL of 50uM hemin complete media was added to the cells for 32h before cells were harvested for analysis. The siRNA sequences used are as follow: Negative Control siC1 from Sigma Universal Negative Control #1 (SIC001); Negative Control siC2 from mit.edu/sirna (GL3 #1008); Negative Control siC3 from mit.edu/sirna (siRL3 #1001); siMAP3K7#1 from Sigma, NM_003188 SASI_Hs01_00234777 CCTGAAACCACCAAACTTAdTdT; siMAP3K7#2 from Sigma, NM_003188 SASI_Hs01_00234778 CATGCAACCCAAAGCGCTAdTdT; siMAP3K7#3 from Sigma, NM_003188 SASI_Hs02_00335227 GTGTTTACAGTGTTCCCAAdTdT; siGATA1#1 from Sigma, NM_002049 SASI_Hs01_00092060 GGAUGGUAUUCAGACUCGAdTdT; siGATA1#2 from Sigma, NM_002049 SASI_Hs01_00092059 CCAAGAAGCGCCUGAUUGUdTdT.

### Western blot analysis

Whole-cell lysates of cultured cells or MDS BM MNCs were prepared with standard radioimmunoprecipitation assay buffer in the presence of protease (Sigma, cOmplete) and phosphatase (Sigma, PhosSTOP) inhibitor cocktails. Protein concentration was measured using Bio-Rad Protein Assay. Protein lysates (15-25 ug) were run on Novex Tris-Glycine gels (Invitrogen) and transferred to nitrocellulose membranes (Bio-Rad), followed by immunoblotting with primary and secondary antibodies using standard methods. Bands were measured and quantified using ImageJ (NIH) or ImageQuant (Molecular Dynamics) softwares. All antibodies used were as follow: anti-MAP3K7 (Cell Signaling, 4505S), anti-p38 MAPK (Cell Signaling, 8690T), anti-p-p38 MAPK (T180) (Cell Signaling, 4511S), anti-p-NF-kappaB p65 (Cell Signaling, 3033P), anti-GATA-1 (Cell Signaling, 3535T), anti-p-HSP27 (Cell Signaling, 9709T), anti-p-MAPKAPK-2 (MK2; T334) (Cell Signaling, 9595T), anti-α-Tubulin (Cell Signaling, 3007T), anti-p-SAPK/JNK (T183/Y185) (Cell Signaling, 9255S), anti-p-ERK1/2 (Bethyl, A303-608A-T), anti-K700E SF3B1 (Bethyl, custom made), anti-SF3B1 (Bethyl, A300-996A), anti-beta-Actin (Sigma, A3853), anti-GAPDH (Sigma, G9545), and anti-HA (Sigma, H3663)

### RT-PCR and qPCR

One microgram of RNA was used for reverse transcription using Invitrogen’s SuperScript III First-Strand kit with the provided oligo-dT (ThermoFisher, cat # 18080-051). For qPCR, transcript levels were measured using SYBR Green qPCR (Applied Biosystems, cat # 4367659). Relative RNA levels in whole-cell lysates were determined by normalizing expression levels of the target genes to expression levels of GAPDH. For RT-PCR, PCR was carried out using EconoTaq PLUS GREEN 2x master mixes (Lucigen) and primers for target genes and control housekeeping genes. RT-PCR primers used are the following: SF3B1 5’-gctgtgtgcaaaagcaagaa-3’ and 5’-agccaaaccctttcctctgt-3’; MAP3K7 5’-GTGAGATGATCGAAGCCCCT-3’ and 5’-CAGCACCATGCAGCACATTA-3’. qPCR primers used are as follow: GAPDH 5’-CTGACTTCAACAGCGACACC-3’ and 5’-CCCTGTTGCTGTAGCCAAAT-3’; LDB1 5’-TGTCACGCCACAAGACCTAC-3’ and 5’-GGAGAGGGCGAAGGTGCTA-3’; HSPA5 5’-GAAAGAAGGTTACCCATGCAGTTG-3’ and 5’-CATTTAGGCCAGCAATAGTTCCAG-3’; c-MYC 5’ -CGTCTCCACACATCAGAGCACAA-3’ and 5’-GCAGCAGGATAGTCCTT-3’; c-MYB 5’-GAAAGCGTCACTTGGGGAAAA-3’ and 5’-TGTTCGATTCGGGAGATAATTGG-3’; and EKLF1 5’-ATGACTTCCTCAAGTGGTGG-3’ and 5’-CTCTCATCGTCCTCTTCCTC-3’.

### Erythroid differentiation of K562 cells

Logarithmically growing K562 (30,000-50,000/mL) cells were treated with a final concentration of 30uM, 40uM, or 50uM hemin (Sigma, cat # 51280-1G) in IMDM media supplemented with 10% FBS for 3d to 4d as previously described^43^. The cells were cultured in a 37°C, 5% CO_2_ incubator.

### Purification, erythroid induction culture, and transduction of human CD34+ cells

Human CD34+ cells were purified from adult peripheral whole blood (LeukoPaks) by positive selection using microbeads magnetic cell separation system (Miltenyi Biotec, Cat # 130-100-453), according to the manufacturer’s instructions. The purity of the isolated CD34+ cells was greater than 95%. CD34+ cells were induced to erythroid differentiation using three phase-culture media cocktails, containing various amount of erythropoietin, as previously described^45^. Transduction of human CD34+ cells with viruses was performed using a method we previously published^44^. The cells were cultured in a 37°C, 5% CO_2_ incubator.

### Erythroblast staging analysis

Healthy and MDS BM aspirates were analyzed for the four erythroblast stages (proerythroblast, basophilic normoblast, polychromatophilic normoblast, and orthochromatic normoblast) using a previously published method^45^. Briefly, BM aspirates were separated on a Ficoll (GE Healthcare) density gradient, and the purified mononuclear cells were incubated with CD45 microbeads (STEMCELL Technologies) for negative selection according to the manufacturer’s instructions. CD45 negative cells were stained with antibodies to detect three erythroid surface markers (GYPA, Integrin Alpha4, Band 3), then flow cytometric analyzed, and quantified as previously described^45^.

### Fluorescence-activated cell sorting of cultured and CD34+ cells

Cells were washed and resuspended in PBS containing 0.5% BSA. The virally-transduced human CD34+ were sorted for GFP reporter expresssion on a MoFlo high-speed cell sorter (Beckman-Coulter) located at the New York Blood Center, NY. Transduced cultured cells were sorted for GFP on an Influx cell sorter (BD Biosciences) or BD FACSAriaII sorter (BD Biosciences) at Columbia University Irving Medical Center.

### Flow cytometric analysis

Cells were washed with PBS containing 0.5% bovine serum albumin (BSA). After blocking with 0.4% human AB serum for 10 minutes on ice, cells were stained with fluorochrome-conjugated antibodies in PBS/0.5% BSA for 15 minutes at 4°C. Cells were washed once with PBS/0.5% BSA before analysis. When Annexin V analysis was performed, cells were washed with 1x Annexin V binding buffer (Biolegend, Cat #422201) instead. Cells were subsequently stained with Fluorochrome-conjugated Annexin V in 1x Annexin V binding buffer according to manufacturer’s protocol. Cells were immediately analyzed on a FACSCalibur (Becton Dickinson) or LSRFortessa (BD Biosciences) cell analyzer. FACS files were analyzed using FlowJo software (FlowJo, LLC). FACS antibodies used were the following: anti-human CD235a (GYPA) PE (Biolegend, 349106), anti-AnnexinV APC (Biolegend, 640941), anti-CD45 APC-Cy7 (Biolegend, 304014), anti-AnnexnV Pacific Blue (Biolegend, 640917), anti-Ki67 AF488 (Biolegend, 151204), 7AAD (BD Pharmingen, 559925), anti-alpha4 integrin/CD49d PE (Miltenyi Biotec, 130-118-548), anti-CD235a-PC7 (PE-Cy7) (Beckman Coulter, A71564), and anti-Band 3 APC (Xiuli An’s lab made^26^).

### Intracellular Ki67 staining

Day 3-hemin-treated cells were washed and stained with GYPA-PE surface marker as described above. Cells were then washed, fixed and stained for intracellular Ki67 following BD Pharmingen’s procedure for FITC BrdU Flow Kit (Cat# 559619) with two modifications: DNase treatment step was omitted and Ki67-AlexaFluor488 (Biolegend #350507) was used instead of anti-BrdU FITC.

### Identification of novel cryptic 3’splice site usage

In order to annotate novel 3’ss that are not present in current datasets, we adopted the splice junction read output by STAR alignment^45^. We adopted the pipeline from DeBoever^16^ for novel splice junction identification and usage, as well as identification of associated canonical 3’ss for cryptic 3’ss. To this end, RNASeq .fastq files were aligned using a splice junction database from DeBoever^16^. Counts of junction reads from SJ.out.tab file (STAR output) were merged into a unique matrix, with each row indicates one splice junction and column for each sample. We further filtered out junctions with a read coverage of <20 summed across all samples. DEXeq was then applied to detect significantly differential mis-spliced 3’ss using the code from DeBoever et al (https://github.com/cdeboever3/deboever-sf3b1-2015). A bar plot of log_2_ distance in base pair was used to demonstrate cryptic 3’ splicing pattern on significantly differentially used novel 3’ss.

### Gene ontology analysis

To determine gene ontology pathways that could potentially be affected by abnormal splicing in *SF3B1* mutant cells. To this end, we first performed standard enrichment analysis using GO David online tools. The input is a list of genes associated with significant aberrant splicing events between mutant and wild-type by satisfying: 1) significantly more frequently used in mutant with *q*-value <0.05; and 2) associated with a distance <50 bp upstream of the canonical 3’ss.

### RNAseq coverage plots of cryptic 3’ss visualization

We used Integrative Genomics Viewer (IGV)^46^ to visualize usage of cryptic 3’ss in *SF3B1* mutant and wild-type cells. For comparisons of different *SF3B1* hotspot mutations, we downloaded MDS RNASeq data from NCBI Sequence Read Archive under accession number GSE85712^18^ and GSE63569^52^. We also downloaded RNASeq data of K562 CRISPR cells with *SRSF2* mutation (GSE128805,^40^) or *U2AF1* mutation (our unpublished data).

### RNAseq expression analysis

To generate mRNA expression matrix for transcriptome analysis, we used featureCounts^47^ from package ‘Subread’ to call read counts from STAR re-aligned bam files. Genes with low read depths across the cohort are removed. Then, read counts were transformed into RPKM values, followed by log2 transformation, and quantile normalized on the sample level.

### Statistical Analysis

Data are expressed as mean ± standard deviation. Comparisons were analyzed by using Student 2-tailed paired with equal variance t-tests. Differences were considered significant at p <0.05.

### Data Availability and Coding Availability

RNA sequencing data reported in this paper will be uploaded to Gene Expression Omnibus (GEO) database upon paper acceptance.

## SUPPLEMENTAL INFORMATION

### SUPPLEMENTAL FIGURE LEGENDS

**Figure S1.**
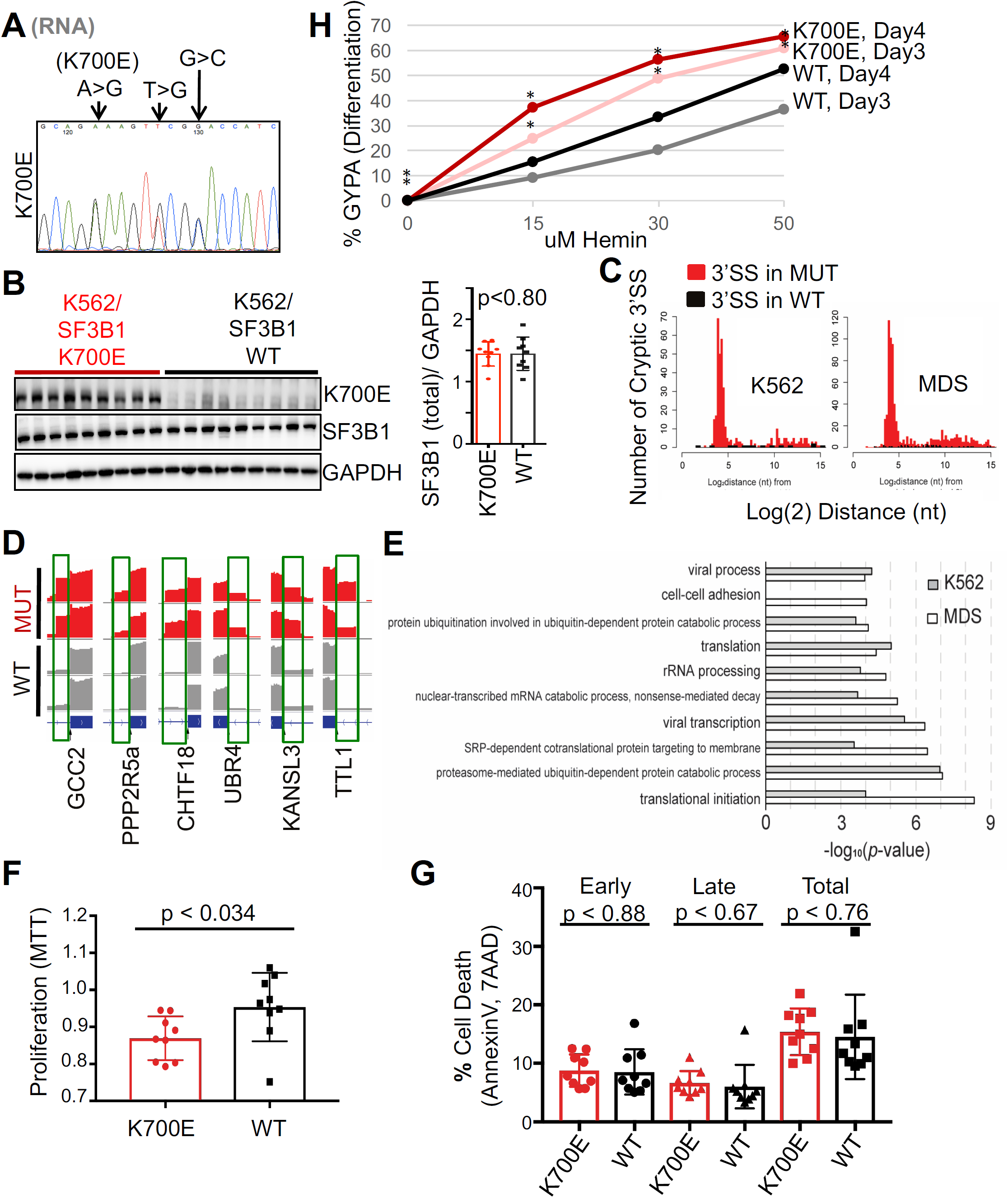
Characteristics of K562/SF3B1 K700E and WT CRISPR clones. (A) DNA chromatogram of RT-PCR products from a representative K562/SF3B1 mutant clone, demonstrating the stable expression of K700E and two silent mutations in *SF3B1* mRNA. (B) Left panel, representative western blot images showing expression of total SF3B1, K700E mutant SF3B1, and loading control GAPDH in the nine independent K700E mutant and nine independent WT K562/SF3B1 clones. Right panel, bar graph displaying the results of ImageJ-quantified, GAPDH-normalized total SF3B1 band intensity and p-value from t-test. (C) Image showing Log^2^ distance in nucleotides from cryptic 3’ss to corresponding canonical 3’ss from K562 cells (left panel) and MDS patients (right panel). (D) RNA-seq coverage plots of some examples of cryptic 3’ss-containing transcripts in two K562/SF3B1 K700E and two WT clones: (from left to right) *GCC2, PPP2R5A, CHTF18, UBR4, KANSL3* and *TTI1*. (E) Image displaying comparison of the top 10 enriched GO pathways determined from misspliced transcripts from SF3B1 mutant K562 cells (grey bar) and SF3B1 mutant MDS patients (white bar). Representative bar graphs quantifying (F) the relative proliferative capacity as measured by MTT assay (n=3 independent experiments) and (G) cell death as measured by FACS using AnnexinV and 7AAD (n=3 independent experiments) in SF3B1 mutant and WT K562 cells under normal growth conditions. p-values from t-tests are shown. (H) Bar graph showing the normalized percent GYPA of Fig. 1E to 0 uM hemin. *, p < 0.05.

**Figure S2.**
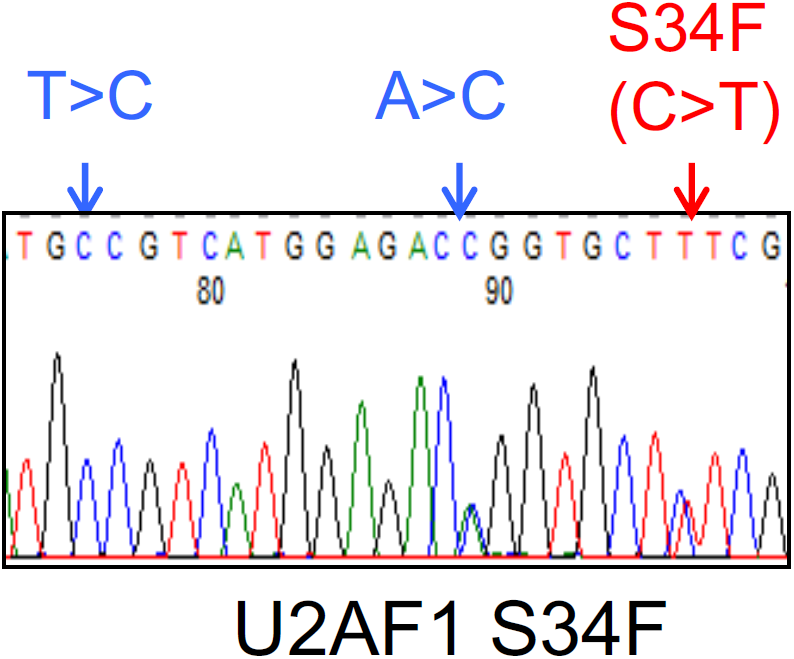
Introduction of the hotspot *U2AF1* S34F mutation into K562 cells using CRISPR/Cas9. DNA chromatogram of representative K562/U2AF1 mutant clone, showing hotspot S34F and two silent mutations in the *U2AF1* gene by CRISPR.

**Figure S3.**
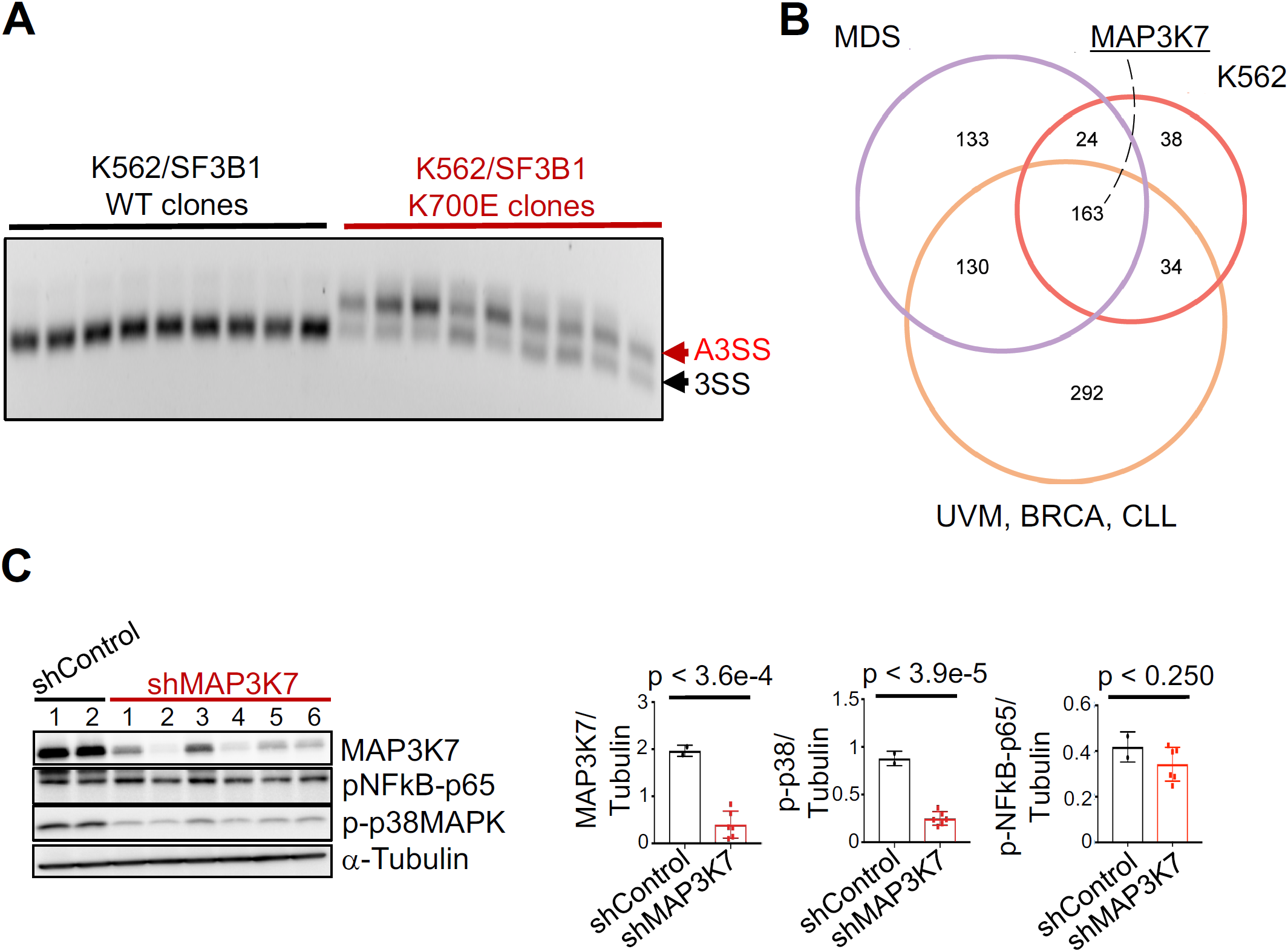
*MAP3K7* is misspliced in mutant *SF3B1* cells from K562, MDS, and other cancers. (A) Image of RT-PCR gel showing 3’ss usage in mutant K562/SF3B1 cells. (B) Venn diagram showing the numbers of differentially spliced transcripts in mutant *SF3B1* samples from MDS, K562/SF3B1 and other cancers: uveal melanoma (UVM), breast carcinoma (BRCA), and chronic lymphocytic leukemia (CLL) (gene list of other cancers from Deboever et al., 2015). (C) Images (left) of representative western blot analysis of GFP-sorted, shRNA-mediated MAP3K7 KD (two different negative control and six different MAP3K7 shRNAs) in parental K562 cells, and bar graphs (right) displaying the results of ImageJ-quantified, α-Tubulin-normalized phospho-NFkBp65, MAP3K7, and p-p38 band intensities in western blot analysis are shown. p-values from t-tests are labelled on all bar graphs. n= 2 independent experiments.

**Figure S4.**
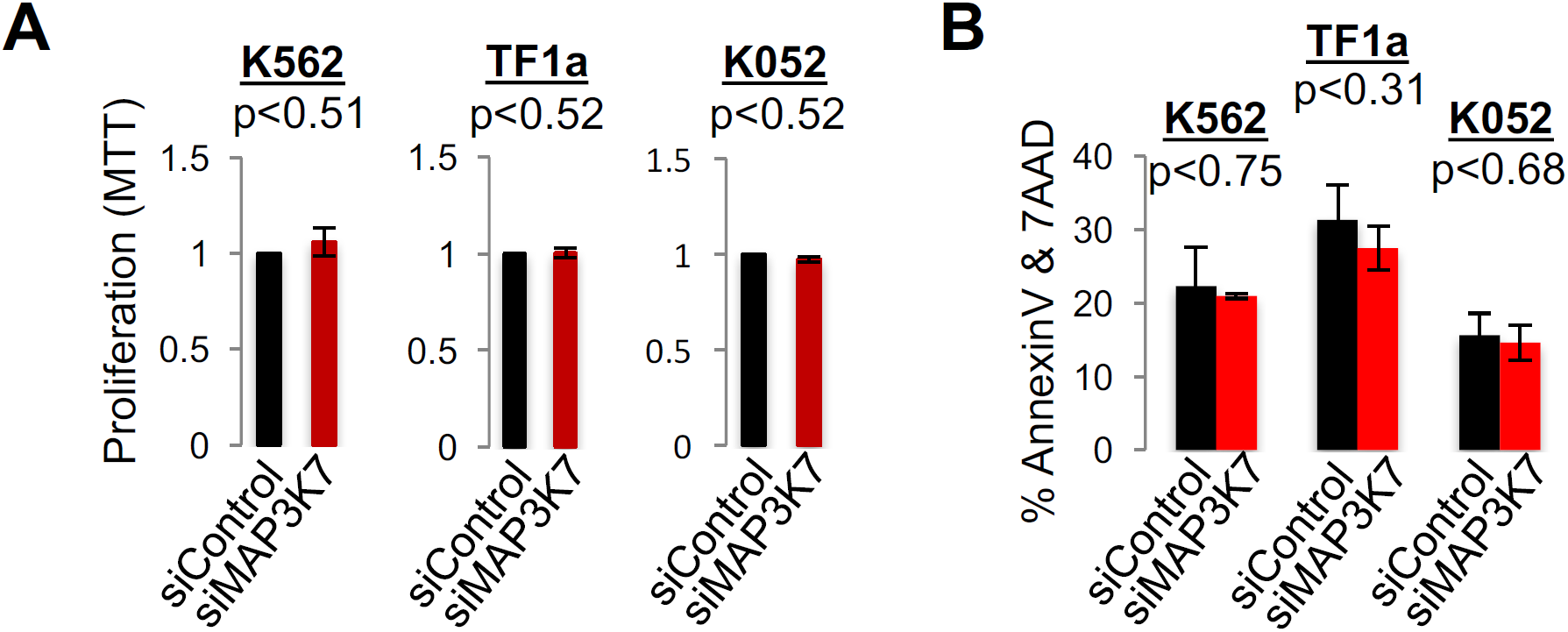
Knockdown of *MAP3K7* in K562, TF1a, and K052 cells has no effects on proliferation and cell death under normal growth conditions but leads to increased erythroid development and erythroid cell death. Bar graphs quantifying (A) proliferation via MTT assay and (B) cell death via AnnexinV and 7AAD by FACS analysis of K562, TF1a, and K052 cells at 48h post-transfection with control (siControl #1) or MAP3K7 (siMAP3K7 #2) siRNAs under normal growth conditions. p-values from t-tests are shown. n=3 independent experiments.

**Figure S5.**
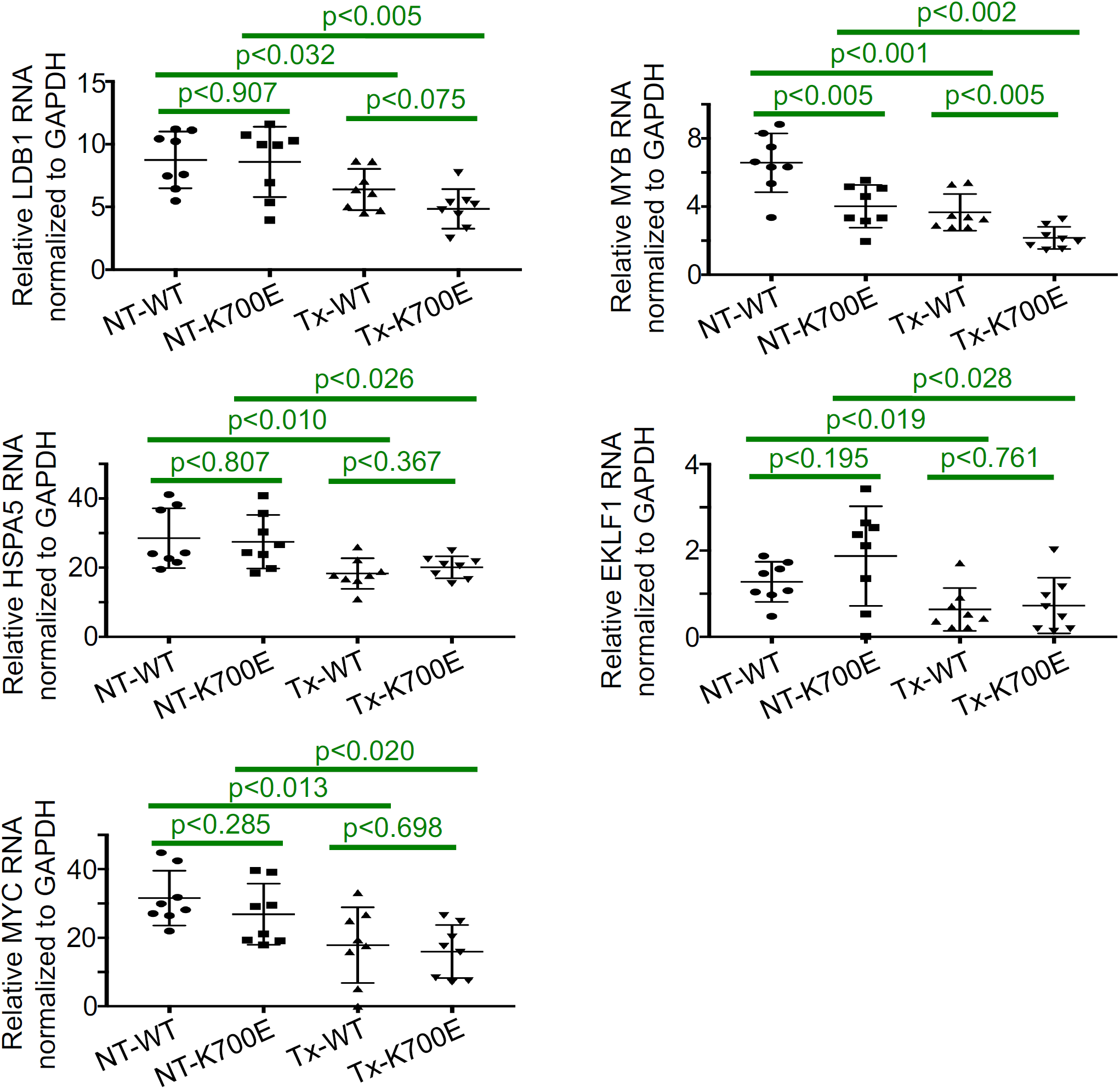
Expression of several major transcription factors in erythroid differentiation of K562/SF3B1 cells. qPCR showing expression of several transcription factors (*LDB1, MYB, HSPA5, EKLF1, MYC*) that are known to play a role in erythroid differentiation. p-values from t-tests are shown.

**Figure S6.**
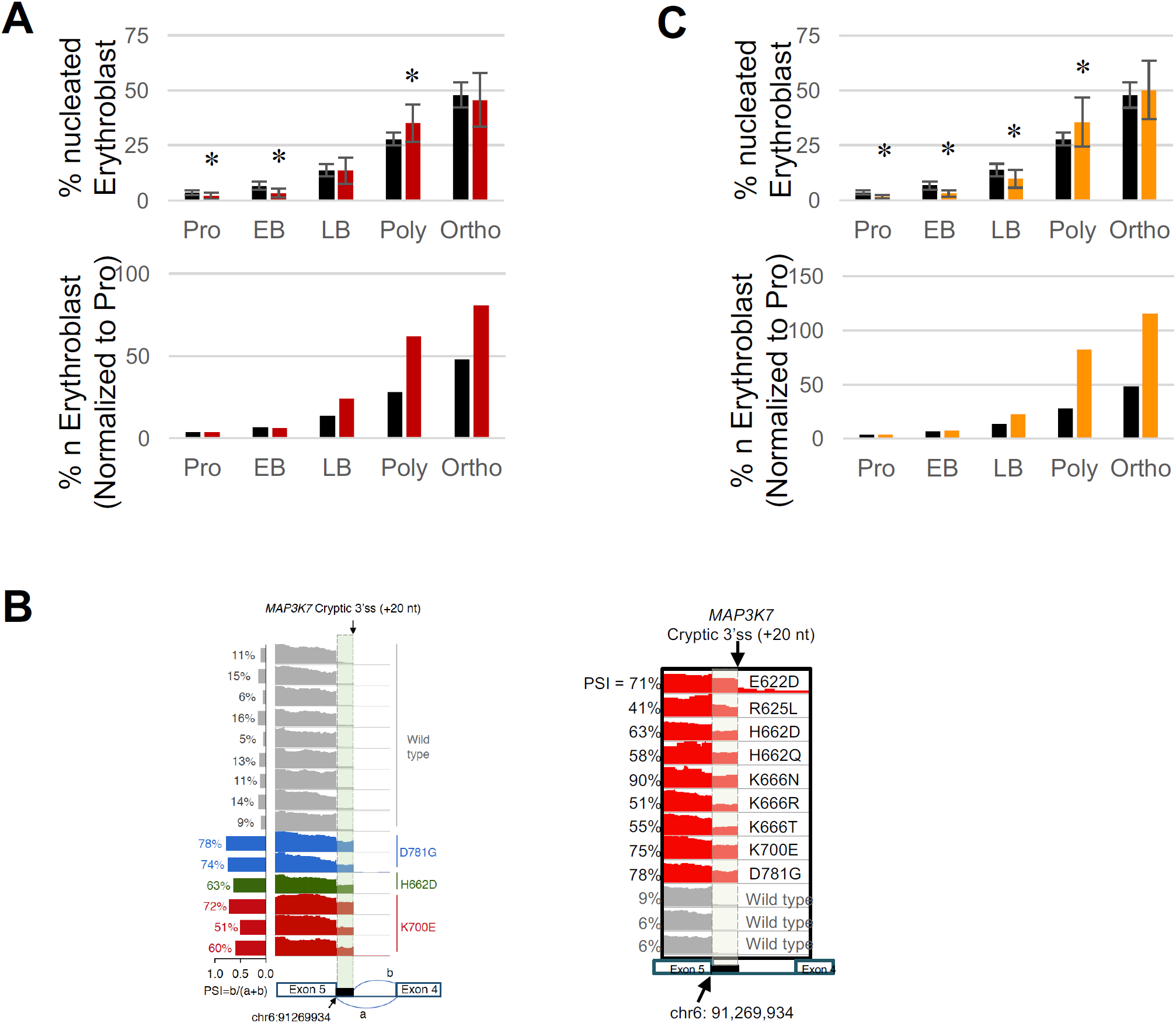
Other MDS SF3B1 mutations, besides K700E, also cause *MAP3K7* missplicing and accelerated erythroid differentiation. Re-plotted MDS erythroblast profile data from Ali et al. (2019) using only normal healthy samples (n=16) and MDS samples (A) with K700E (n=15) and (C) with non-K700E SF3B1 mutations (n=14). The percentages of nucleated erythroid cells in each erythroblast stage are illustrated in the bar graphs. *, p < 0.05. Normalized-to-proerythroblast (Pro)-staged bar graphs are also shown for comparison. Pro, proethroblasts; EB, early basophilic erythroblasts; LB, late basophilic erythroblasts; Poly, polychromatic erythroblasts; and Ortho, orthochromatic erythroblasts. (B) RNA-seq coverage plots of *MAP3K7* cryptic 3’ss in MDS patients with different *SF3B1* hotspot mutations (MDS RNASeq data downloaded from Zhang et al. 2019 (left panel) and from both Dolatshad et al. 2015 and Obeng et al. 2016 (right panel)) and with no *SF3B1* mutations (wild-type).

**Figure S7.**
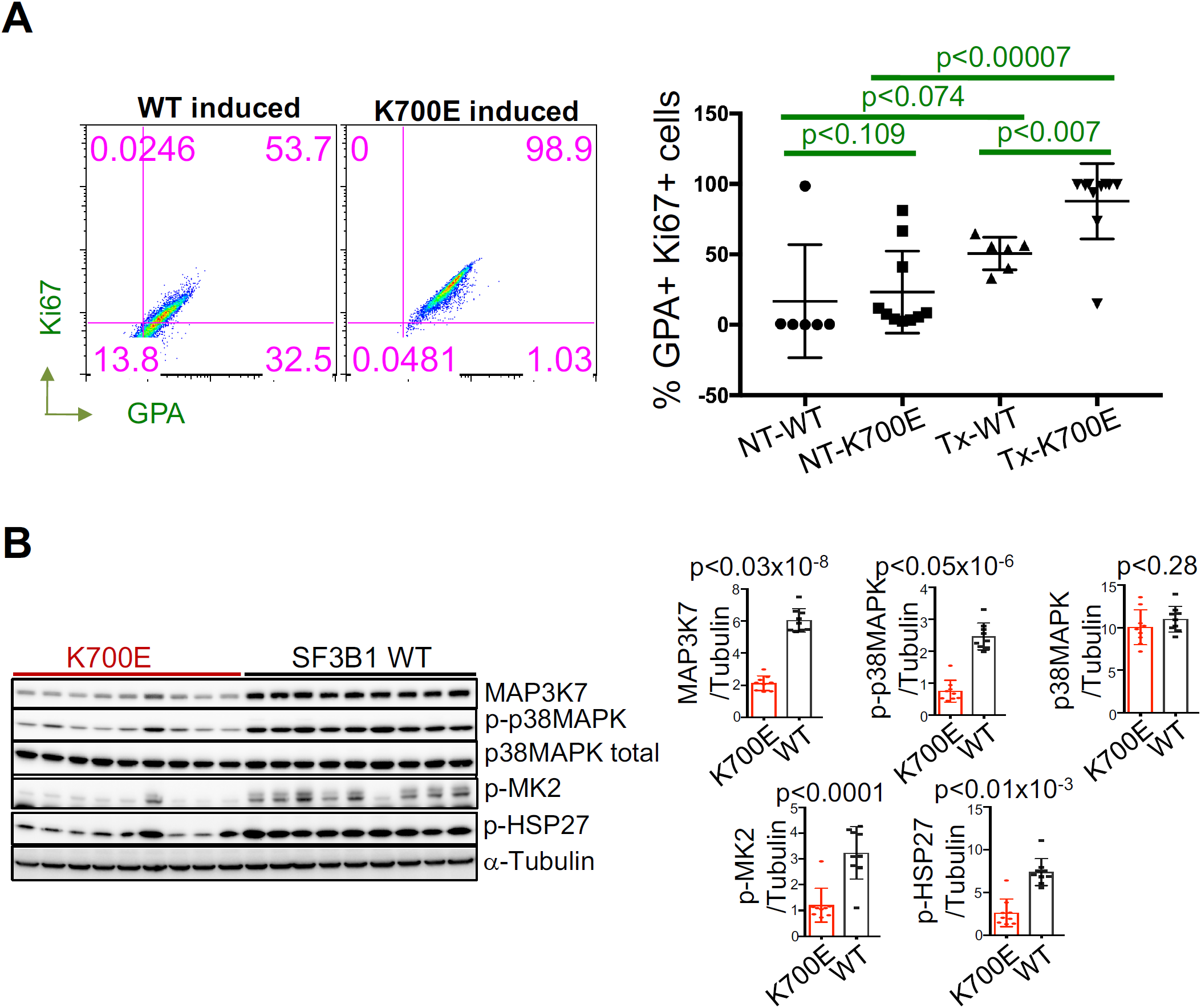
Mutant SF3B1 cells display increased proliferative capacity and deactivation of p38 MAPK downstream effectors, MAPKAPK2 (MK2) and HSP27. (A) Expression of proliferative marker Ki67 in mutant and WT K562/SF3B1 erythroid cells. FACS analysis of Ki67 vs. GYPA expression in representative 50 uM hemin-induced day 3 K700E and WT cells. Results of multiple K700E and WT clones are summarized graphically. p-values from t-tests are shown. n=2 independent experiments. (B) Representative western blot images showing expression of p-MK2 and p-HSP27 in mutant and WT K562/SF3B1 clones. Bar graphs display the results of ImageJ-quantified, α-Tubulin-normalized protein band intensities and p-values from t-tests. n=3 independent experiments.

### SUPPLEMENTAL TABLE TITLES

**Table S1. Cryptic 3’ Splice Sites Differentially Used by Mutant 700E vs WT SF3B1 K562 Cells, Related to Figure 1B.**

**Table S2. Cryptic 3’ Splice Sites Differentially Used by Mutant vs WT SF3B1 MDS Cells, Related to Figure 1B.**

